# Fluorescence visualization of deep-buried hollow organs

**DOI:** 10.1101/2022.01.07.475462

**Authors:** Zhe Feng, Yuanyuan Li, Siyi Chen, Xiaoming Yu, Yanyun Ying, Junyan Zheng, Tianxiang Wu, Jin Li, Xiaoxiao Fan, Dan Zhang, Jun Qian

## Abstract

High-definition fluorescence imaging of deep-buried organs is still challenging. Here, we develop bright fluorophores emitting to 1700 nm by enhancing electron donating ability and reducing donor-acceptor distance. In parallel, the heavy water functions as the solvent of the delicately designed fluorophores, effectively reducing the fluorescent signal loss caused by the absorption by water. The near-infrared-II (NIR-II, 900-1880 nm) emission is eventually recovered and extended beyond 1400 nm. Compared with the spectral range beyond 1500 nm, the one beyond 1400 nm gives a more accurate fluorescence visualization of the hollow organs, owing to the absorption-induced scattering suppression. In addition, the intraluminal lesions containing much water are simultaneously negatively stained, leading to a stark contrast for precise diagnosis. Eventually, the intraluminally perfused fluorescent probes are excreted from mice and thus no obvious side effects emerge. This general method can provide new avenues for future biomedical imaging of deep and highly scattering tissues.

## Introduction

The second near-infrared (NIR-II, 900-1880 nm) window gives a promising fix for deep-penetration biological imaging^1-6^. The vessels, lymph, tumor, etc., have been successfully visualized intravitally by virtue of NIR-II fluorescence imaging^7-16^. Besides the reduced photon scattering, the rising light absorption by water further restrain the scattering disturbance in tissues during NIR-II intravital imaging^17, 18^. The image contrast is thus improved and the NIR-IIx+NIR-IIb region (1400-1700 nm) is promoted as one highly-potential imaging window^17^. However, it must be admitted that there is still a shortage of bright probes that could achieve the long-wavelength emission and enough integrated photoluminescence intensity. On the other hand, aqueous dispersion of the exogenous probes is a major requisite for most biomedical researches, thus breeding many advanced hydration techniques for liposoluble emitters. Compared with the first near-infrared region (NIR-I, 760-900 nm), the long-wavelength-emitting fluorophores are confronted with non-negligible depletion problems in the NIR-II window, especially beyond 1400 nm^19^.

To improve the brightness of collected signals, enhancing the absolute photoluminescence of the fluorophores by molecular design to resist the absorption loss in the biological environment is extremely essential. In this work, we propose a novel molecular design strategy of donor engineering to enhance the emission in the long-wavelength spectral regions. Correspondingly, we present a general method to suppress the imaging background and recover the signals with full capitalization on the water in the tissue and the heavy water of solvent.

The good biocompatibility of organic fluorophores could facilitate their biomedical applications^20-26^. Highly bright fluorophores are generally designed as twisted structures full of molecular rotors to suppress the undesirable intermolecular π-π interactions in the aggregated state^27-30^. On the basis of this principle, a series of highly bright NIR-II molecules with absorption and emission peaks around 700 and 1000 nm, respectively, have been successfully achieved^31-33^. However, NIR-II dyes with longer spectral responses and brighter emission are still challenging to obtain owing to the decrease of emission efficiency with an increase of wavelength. To solve this issue, we enhanced donor-acceptor (D-A) interactions through the reduction of D-A distance based on the guideline of “backbone distortion and molecular rotors” (see Fig. 1a), instead of simply increasing the conjugation length. The absorption and emission are driven to the longer-wavelength range and the high photoluminescence intensity is simultaneously maintained. Particularly, the as-synthesized 2FT-oCB molecule showed a strong absorption peak at 828 nm (molar extinction coefficient, *ε* of 2.3×10^4^ M^-1^ cm^-1^) and an emission peak at 1145 nm extending to 1700 nm.

**Figure 1.**
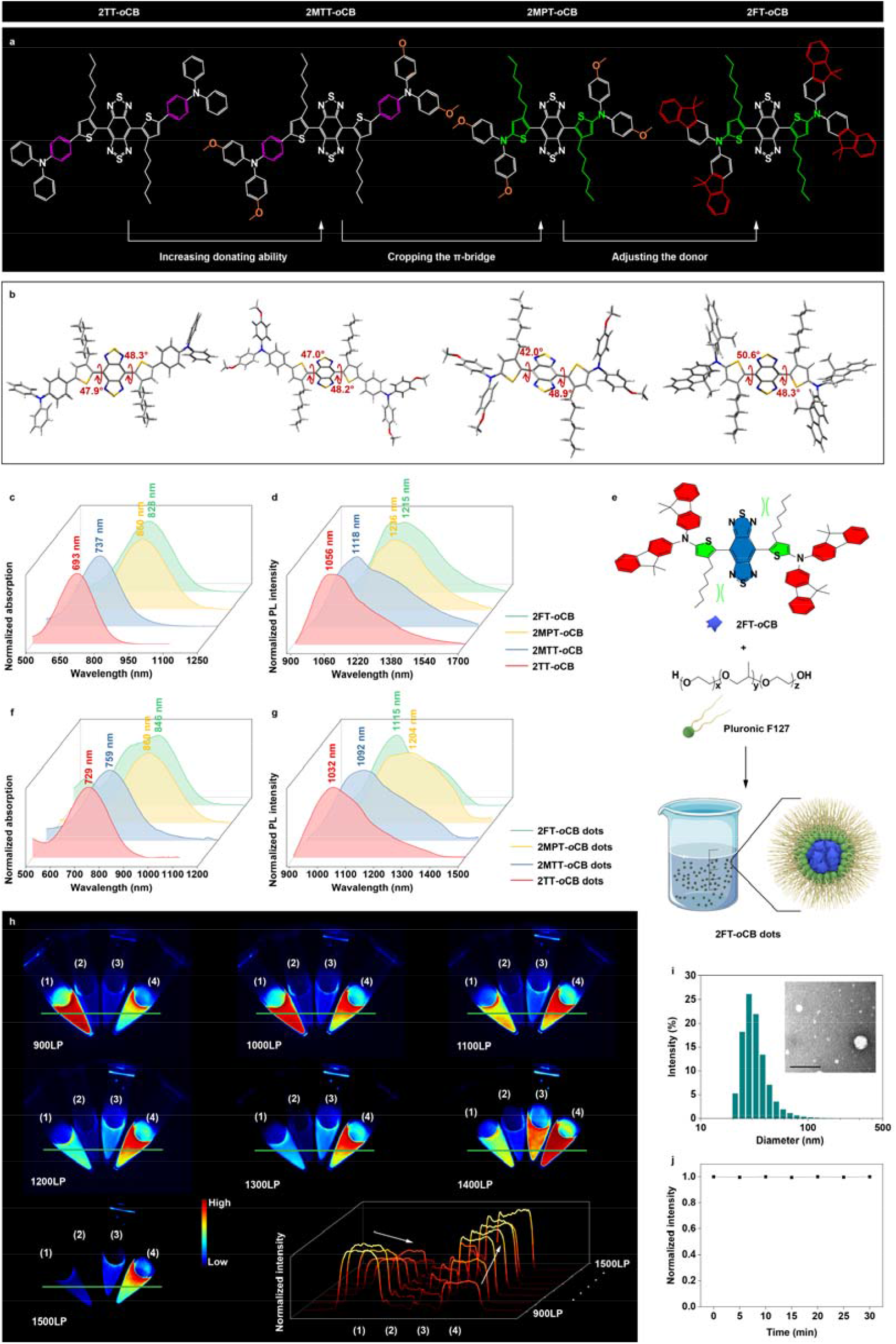
The molecular engineering of NIR-II fluorophores with bright and wavelength-extended emission. **(a)** The molecular structures and **(b)** the optimized S_0_ geometries of the four molecules (2TT-*o*CB, 2MTT-*o*CB, 2MPT-*o*CB and 2FT-*o*CB). **(c)** The normalized absorption and **(d)** the normalized PL spectra of the four molecules dissolved in tetrahydrofuran (THF). **(e)** Schematic illustration of the fabrication of fluorescent dots. **(f)** The normalized absorption and **(g)** the normalized PL spectra of the four kinds of AIE dots dispersed in water. **(h)** The normalized fluorescence images of tubes containing the aqueous dispersion of four fluorescent dots with the same concentration and the normalized cross-sectional fluorescence intensity profiles along the green lines. (1), 2TT-*o*CB dots; (2), 2MTT-*o*CB dots; (3), 2MPT-*o*CB dots; (4), 2FT-*o*CB dots. **(i)** The DLS results of the 2FT-*o*CB dots. The insert is the representative TEM image of the 2FT-*o*CB dots negatively stained with uranyl acetate. Scale bar, 100 nm. **(j)** The photostability test of the 2FT-*o*CB dots under continuous irradiation.

Furthermore, a parallel efficient strategy was carried out to transfer the solvent of fluorescent probes from natural water (hydrogen oxide) to heavy water (deuterium oxide) to recover the bright NIR-II emission, since the stable deuteration of water could efficiently red-shift the light absorption with no radiological concerns introduced^34^. To avoid the disturbance of deuterium oxide to the internal biological environment, we tapped the visualization potential of deep-buried hollow organs whose deciphering is always obstructed by the skin, muscle, fat, and other upper organs. To facilitate the high-performance hollow organ imaging, we collected the emission of NIR-IIx (1400-1500 nm) extra with larger light absorption to further suppress the imaging background, diverging from traditional collection beyond 1500 nm (NIR-IIb window) where only the photon scattering is considered. This method stores a high potential for clinical translation considering the intraluminal perfusion instead of direct injection into the circulatory system excludes the major safety concerns of exogenous agents. We believe our findings presented in this work will be transformational for further biomedical researches and future clinical applications.

## Results

### Molecular design and fluorophore synthesis

The chemical structures of 2TT-*o*CB, 2MTT-*o*CB, 2MPT-*o*CB and 2FT-*o*CB are shown in Fig. 1a. The synthetic route and structural characterization of the above molecules associated with key intermediates are displayed in Supplementary Schemes 1-4 and Supplementary Figs. 1-15. The strong electron-withdrawing ability of the central BBTD core can drive the absorption/emission to the long-wavelength range when coupled with electron donors. Ortho-hexyl-substituted thiophene can effectively distort the thiophene-BBTD-thiophene backbone to suppress the intermolecular π-π interactions. Importantly, through molecular rotor engineering, four dyes of 2TT-*o*CB, 2MTT-*o*CB, 2MPT-*o*CB and 2FT-*o*CB with triphenylamine, methoxy-triphenylamine, methoxy-diphenylamine, and fluorene-based diamine, respectively, at the periphery were designed. From 2TT-*o*CB to 2MTT-*o*CB, the introduction of methoxy electronic donating group increases D-A interactions, as reported by previous contributions which focus on the increasing the numbers of donors^35-37^ or D-A units^38-40^. These molecular design strategies open up new avenues to develop high-performance fluorescent molecules. However, the improvement of the conjugation length brings not only redshifted spectral response but declined quantum yield. In this study, we propose a strategy of cropping the π-bridge to enhance the fluorescence efficiency of NIR fluorophores. From 2MTT-*o*CB to 2MPT-*o*CB, the removal of the phenyl unit in triphenylamine may significantly enhance the D-A interactions. Such transformation further twists the molecular structure, effectively avoiding the π-π stacking. Next, 2FT-*o*CB with electronic donating fluorine unit is designed to regulate the D-A interactions. The introduction of the planar blocks aims to improve the molar extinction coefficient. To confirm the chemical configuration, density functional theory (DFT) calculations were carried out using Gaussian 09 program. The optimized conformations show that all the molecules adopt twisted architectures with dihedral angles of ∼50° (Fig. 1b), confirming the steric hindrance between BBTD and *ortho*-positioned alkyl chains. Moreover, the low energy gaps of ∼1.3 eV suggest strong absorption in the NIR biological window (Supplementary Fig. 16).

The photophysical properties of dyes were studied by UV-vis-NIR and photoluminescence (PL) spectroscopy. The effect of enhancing D-A interactions on the dyes was investigated in the tetrahydrofuran (THF) solution. 2TT-*o*CB, 2MTT-*o*CB, 2MPT-*o*CB and 2FT-*o*CB show generally increased maximal absorption wavelength at 693, 737, 860 and 828 nm, respectively (Fig. 1c), with respective *ε* of 0.4×10^4^, 1.0×10^4^, 1.4×10^4^, and 2.2×10^4^ M^-1^ cm^-1^ at the biological window of 793 nm (Supplementary Fig. 17). This result indicates that increasing D-A interactions including the three steps in Fig. 1a can not only redshift the maximal absorption wavelength but also boost the absorption intensity. Notably, such a strong *ε* (2.2×10^4^ M^-1^ cm^-1^) of 2FT-*o*CB at 793 nm is favorable for excitation light to penetrate deep tissues with low photodamage. PL spectra suggest that 2TT-*o*CB, 2MTT-*o*CB, 2MPT-*o*CB, and 2FT-*o*CB display emission peaks at 1056, 1118, 1236, and 1215 nm in THF, respectively, which can be applied as NIR-II emitters for deep tissue visualization (Fig. 1d). The redshifted absorption and emission spectra indicate that both improving the donating ability and reducing the length of π-bridge can efficiently enhance the D-A interaction. Generally, molecular dyes always suffer fluorescence quenching dispersed from good (organic solvents) to poor solvent (water). Modifications of molecular structure for water solubility enhancement or aggregation avoidance are still challenging to keep strong emission at long wavelengths in the NIR-II region. Aggregation-induced emission effect could be an efficient strategy in parallel to manufacture bright long-emitting organic fluorophores^41-43^. To further investigate the fluorescence properties in the aggregated state, the PL spectra of dyes in the THF/water mixture with different water fractions (*f*_w_) were carried out. As shown in Supplementary Fig. 18, the PL intensity of dyes decrease first from *f*_w_ = 0 to *f*_w_ = 40% due to the formation of dark twisted intramolecular charge transfer state (a dark state that can quench the fluorescence), while increase significantly with an increase of *f*_w_ from 40% to 90% owing to the trigger of aggregation-induced emission effect via the mechanism of restriction of intramolecular motion by aggregation formation.

To further decipher the fluorescence properties, we encapsulated the dyes into dots by the nanoprecipitation method using biocompatible surfactant Pluronic F127 (Fig. 1e). Except 2MPT-*o*CB dots (860 nm), the 2TT-*o*CB, 2MTT-*o*CB, and 2FT-*o*CB dots display obviously redshifted maximal absorption wavelength as 729, 759 and 846 nm, respectively (Fig. 1f), compared with the solution profiles (Fig. 1c). On the other hand, the dots show fluorescence at the NIR-II region at 1032, 1092, 1204, and 1115 nm, respectively (Fig. 1g), slightly blue-shifted than their solution state (Fig. 1d). The excellent absorption (> 840 nm) and emission (> 1100 nm) properties of 2MPT-*o*CB and 2FT-*o*CB dots are particularly favorable for high-quality bioimaging. The fluorescence images of 2TT-*o*CB, 2MTT-*o*CB, 2MPT-*o*CB and 2FT-*o*CB dots with same concentrations were recorded using seven different long-pass (LP) filters (900, 1000, 1100, 1200, 1300, 1400 and 1500 nm) (Fig. 1h). Under 900 and 1000 nm LP filters, 2TT-*o*CB dots display the strongest fluorescence signal owing to the high quantum yield (8.4%) in the NIR-II region. Fig. 1h demonstrates that simply increase the donating ability from 2TT-*o*CB to 2MTT-*o*CB induces obvious decline of fluorescence intensity owing to the enhanced D-A interactions. However, by cutting the π-bridge (D-A interaction obviously increased), the wavelength of the maximal absorption/emission of 2MPT-*o*CB is redshifted (see Fig. 1c, d, f and g). Notably, 2MPT-*o*CB dots still maintain strong fluorescence intensity owing to the highly twisted architecture without phenylene π-bridge (see Fig. 1h). With an increase of wavelength of filters from 1100 to 1500 nm, an overwhelming fluorescence signal is observed in 2FT-*o*CB dots decorated with highly twisted skeletons and planar fluorene units, compared with the other three dots. It is noticeable that 2FT-*o*CB dots possess the strongest fluorescence intensity under the 1400 nm LP filter. Therefore, we chose 2FT-*o*CB dots from the designed four fluorophores as the example for the following *in vivo* imaging. As shown in Fig. 1i, the 2FT-*o*CB dots coated by the F-127 possess a homogeneous spherical structure, with diameters around 50□nm measured by transmission electron microscopy (TEM) and dynamic light scattering (DLS). Importantly, the 2FT-*o*CB dots show almost no decline of fluorescence intensity after 30 min of the continuous laser irradiation (see Fig. 1j), which demonstrates their excellent photostability.

### Natural water in the biological tissue improves the imaging quality

The visible-NIR imaging windows are shown in Fig. 2a, which are defined according to the photon scattering and light absorption by tissue. To directly compare the *in vivo* performance in different imaging windows and determine the optimal long-pass detection region of NIR-II fluorescence off-peak imaging, the whole-body vessel imaging in mice was then conducted after intravenous injection of 2FT-*o*CB dots (1 mg/mL, 200 μL 1×PBS solution with normal water). As shown in Supplementary Fig. 19, the images present improved qualities in general, with the rising of the imaging wavelength. Especially, owing to the light absorption, the strong background in the liver induced by the accumulation of 2FT-*o*CB dots was suppressed with 1400-nm long-pass collection, making the vessel network clearly outlined (see Fig. 2b-d). As shown in Fig. 2e, the vessel of interest in the 1400-nm long-pass fluorescence image shows the best SBR of 1.27 while the same vessel possessed the SBRs of 1.07 and 1.16 in 1300-nm long-pass and 1500-nm long-pass images, respectively. The spatial frequency distributions of the images in various regions can be seen in the fast Fourier transform (FFT) results (see Supplementary Fig. 20 and Supplementary Fig. 21a). Similarly, the statistical analysis exhibited a wavelength-related trend in which the intensity of high spatial frequency was positively correlated to the restrained photon scattering approximately. As is known, 980 nm is a typical light absorption peak of water, where the imaging background could be suppressed to some extent. Supplementary Fig. 21c points that the 900-nm long-pass image showed a comparable intensity of high spatial frequency with the 1000-nm long-pass image. Moreover, the spatial-frequency maps of 1300-, 1400-, and 1500-nm long-pass fluorescence images are presented in Fig. 2f-h. Due to the intense light absorption at ∼1450 nm by water, the background arising from the diffused components is highly inhibited, leading that the NIR-IIx+NIR-IIb (1400-1700 nm) image possesses more high spatial frequency even than the NIR-IIb (1500-nm long-pass) image which has been long widely acknowledged with the best quality (Fig. 2i and Supplementary Fig. 21b).

**Figure 2.**
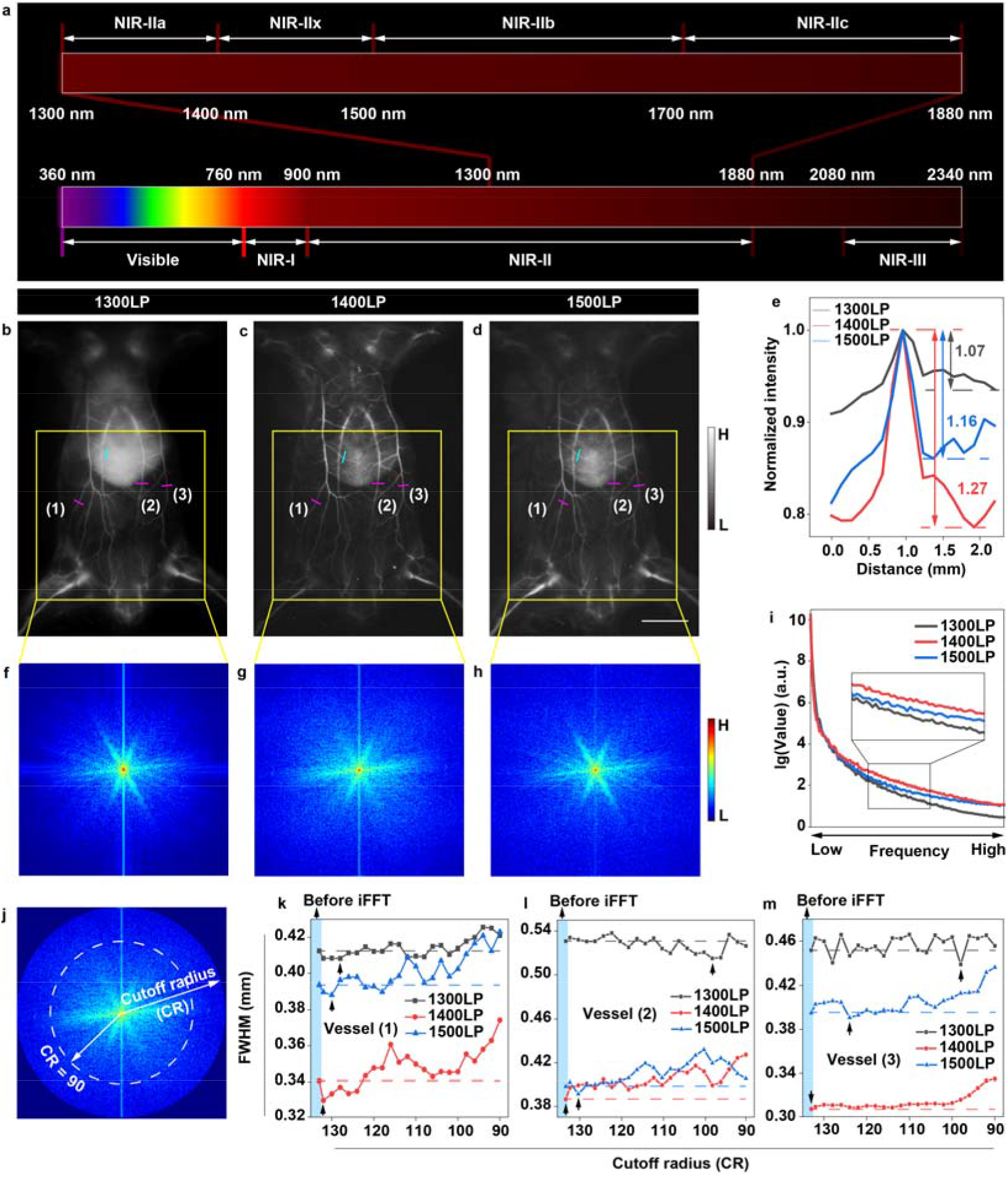
The natural water in the biological tissues optimizes the NIR fluorescence imaging. **(a)** The visible-NIR imaging windows. The **(b)** 1300-, **(c)** 1400- and **(d)** 1500-nm long-pass whole-body fluorescence images in the same mouse after intravenous injection of 2FT-*o*CB dots. Scale bar, 10 mm. **(e)** Cross-sectional fluorescence intensity profiles along the indigo lines of the blood vessel in Fig. 2b-d. The spatial frequency maps after FFT of the **(f)** 1300-, **(g)** 1400- and **(h)** 1500-nm long-pass images. The spatial frequency gradually increases outward from the center of the map, and the color bar indicates the intensity. **(i)** The spatial frequency distribution of Fig. 2f-h. **(j)** The FFT result of the 1400-nm long-pass image where the points in a circumference possessed the same spatial frequency and the cut-off radius represented the cut-off frequency of short-pass filtering. The FWHMs of the **(k)** vessel (1), **(l)** vessel (2), and **(m)** vessel (3) in Fig. 2b-d after short-pass filtering with decreasing cut-off radius, and iFFT. The black arrows pointed to the minimal measured diameters of one vessel in the image.

Random noise is non-negligible interference in the InGaAs-based shortwave infrared (an alternate name of the NIR-II region, which is usually utilized to describe 900-1700 nm in the industrial world) detection. Since the random disturbance usually acts with high spatial frequency in one image, low-pass filtering was used for image denoising. Fig. 2j shows the frequency domain map with the circular distribution of the 1400-nm long-pass image, where the components with a specific spatial frequency are distributed in a circumference with a certain radius. With the decrease of the cut-off radius, high-frequency noise is constantly suppressed. However, detailed information of a picture also corresponds to high-spatial-frequency components which may be wasted through wave filtering without distinction. The diameters of three selected bright vessels in Fig. 2b-d were measured after each inverse fast Fourier transform (iFFT). As shown in Fig. 2k-m, due to the dual reduction of random noise and useful details, each vessel exhibits an optimal cut-off, where the FWHM is minimized. Interestingly, the best FWHM of the vessels in the 1400-nm long-pass image almost emerges without short-pass filtering, which fully reveals that detailed signals in the imaging with 1400-nm long-pass collection account for a considerable proportion in high spatial frequency components, shrinking the disturbance of random noise.

### Heavy water dispersion enhances the NIR-II emission

The absorption by O-H bonds in water results in the light absorption features with the peak wavelengths at ∼980 nm, ∼1200 nm, ∼1450 nm, and ∼1930 nm. In the whole NIR-II region, the absorption at ∼1450 nm is most pronounced. Thus, the growing light attenuation from 1400 nm would inevitably deplete the emission of the fluorophores dispersed in water. The typical absorption spectra of water and heavy water (deuterium oxide) given in Fig. 3a reveal that the fluorescence loss in the NIR-II region induced by solvent could not be ignored anymore, however, the characteristic absorption peak wavelength of water at ∼1450 nm is red-shifted beyond the NIR-II region in heavy water. Accordingly, the dispersion of 2FT-oCB dots in heavy water is considered to be able to recover the emission loss near 1450 nm. After ultrafiltration, the heavy water dispersion was obtained by re-diluting the concentrated 2FT-oCB dots. The PL spectrum of 2FT-*o*CB dots expands in the heavy water dispersion and the peak emission wavelength is red-shifted by 30 nm (see Fig. 3b). Besides, the enlarged spectrum exhibits a highly efficient fluorescence recovery beyond 1400 nm. The emission intensity in the whole NIR-II region of the dots dispersed in heavy water is twice that of dots dispersed in water. Moreover, the emission in heavy water beyond 1400 nm was calculated to be ∼20 times that in water. It would be noteworthy that the PL spectra were measured with the 793 nm laser exciting on the edge of the cuvette, which minimized the absolute fluorescence loss caused by the water. The realistic intensity attenuation would be exacerbated in varying degrees in the different situations, due to the differentiated propagation and assorted compositions. With the IR-26 in the 1,2-dichloroethane chosen as the reference, the quantum yield of 2FT-*o*CB dots beyond 1400 nm can be calculated as 0.11% (see Supplementary Fig. 22).

**Figure 3.**
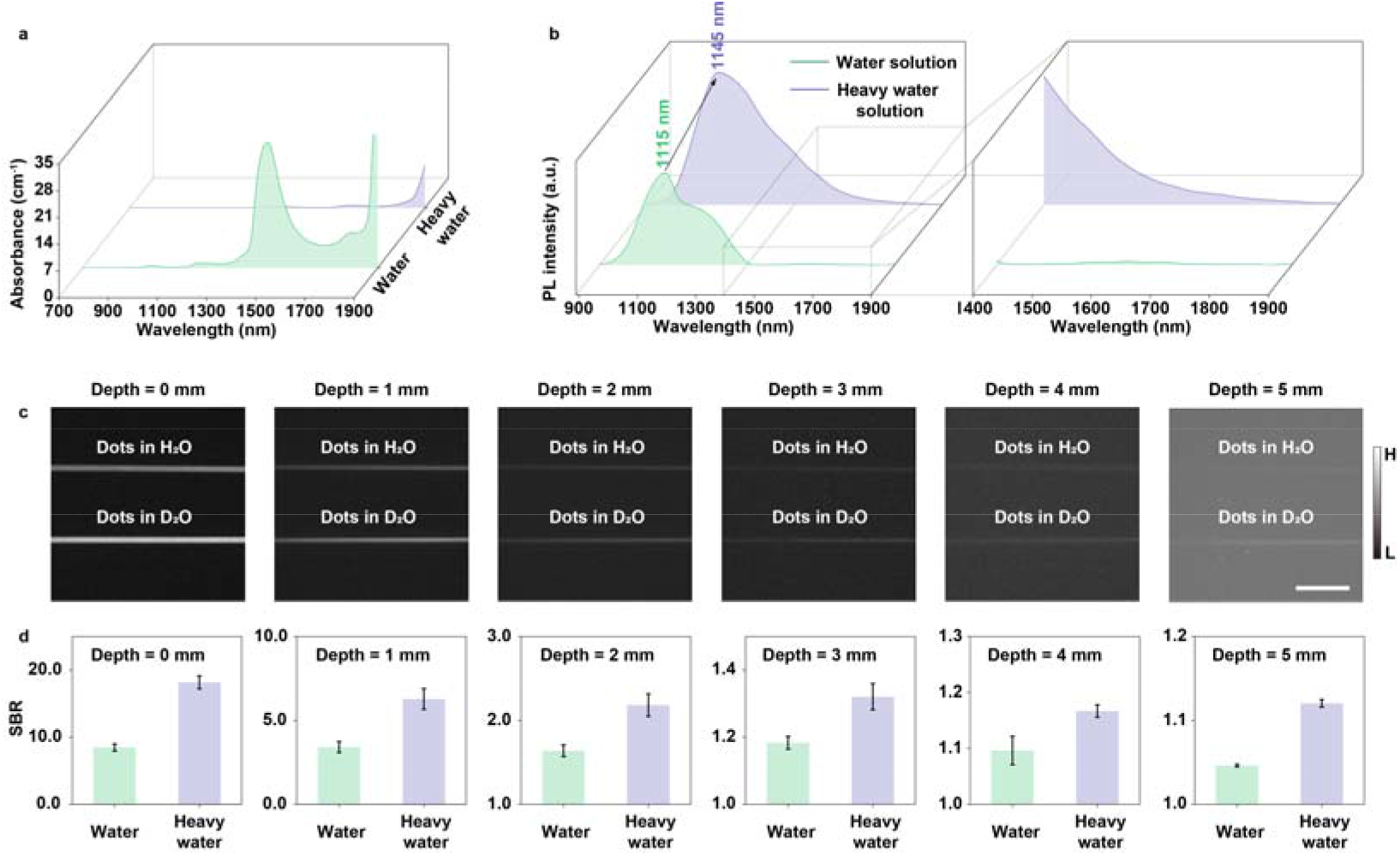
The heavy water dispersion recovers the NIR-II emission of the fluorescent dots. **(a)** The absorbance of water and heavy water in the NIR-I and NIR-II region. **(b)** The PL spectra of 2FT-oCB dots in water and heavy water with the same concentration and the enlarged spectra from 1400 nm to 1900 nm. **(c)** The phantom images of capillaries filled with the hydrogen oxide (top) and deuterium oxide (bottom) dispersion of 2FT-*o*CB dots at depths of 0, 1, 2, 3, 4, and 5 mm in 1% Intralipid^®^ solution with 1400-nm long-pass detection. Scale bar, 5 mm. **(d)** The SBR analyses of the capillaries in Fig. 2c. Error bars indicate s.e.m. (n = 3).

2FT-*o*CB dots possess remarkable emission peaking at ∼1145 nm, which can be recognized as a kind of satisfied fluorescent agent. The Intralipid^®^ phantom study was then conducted to simulate the hindrance of light propagation in biological tissues. As the thickness of 1% Intralipid^®^ solution rose, the images of capillaries (filled with heavy water dispersion of 2FT-oCB dots) beneath become more and more blurred resulting from the increasing photon scattering, as shown in Supplementary Fig. 23a. By and large, the prolonging of detection wavelength can be constructive to ameliorating the imaging results. However, the imaging with 1400-nm long-pass collection, in particular, rather than the 1500-nm long-pass collection (NIR-IIb region) shows the best FWHM, which can be seen in Supplementary Fig. 23b. The seemingly abnormal results can be attributed to the positive contribution of light absorption^17, 18^. In addition, compared with the water dispersion, Fig. 3c-d shows that heavy water dispersion can improve the SBR because of the signal enhancement. The fluorescence recovery via re-dispersing 2FT-oCB dots in heavy water is believed to be an effective and essential approach to the hollow organ visualization, with organ perfusion instead of direct injection into the internal environment.

### High-performance *in vivo* fluorescence cystography and colonography

Fluorescence imaging had long developed a reputation as a technological laggard in deep penetration. The emergence of NIR-II fluorescence imaging brought an infusion of hope to outline the deep details *in vivo* to a certain extent. Ordinarily, it was believed that deeper deciphering required imaging in the longer wavelength region.

Bladder disorders are common as we age, which influence our health in the urinary system. The observation of bladder morphology can assist diagnose the bladder rupture, tumor, and diverticulum to some degree. Near-infrared fluorescence cystography through the skin and muscle was conducted after intravesical instillation of the 2FT-*o*CB dots (1 mg/mL, 20 μL, deuterium oxide dispersion). The non-invasive NIR-II fluorescence image of bladder was set into a green channel, while the bladder image after opening the abdomen was set into a red channel for comparison. Then two images were merged into one and the areas with high consistency were presented in yellow. As shown in Supplementary Fig. 24a-c, the near-infrared emission detection here was progressively red-shifted to restore the original appearance of biostructure as much as possible. The accurate quantification by structural similarity index measure (SSIM) can assist the image quality assessment. As displayed, The SSIM maps are given in Supplementary Fig. 24d in the corresponding imaging window as complementary evaluation for the merged image with quantification. The merged bladder image in 1400-1700 nm shows the best matching, and the presentation exceeds the performance in the NIR-IIb region (see Supplementary Fig. 24c). The mean SSIM indexes in the whole SSIM maps were shown in Supplementary Fig. 24e. With the prolonging of the imaging window, the SSIM index gradually improved and peaked at the map in 1400-1700 nm. The decline of background suppression due to the decreasing light absorption by tissue, and the less NIR-IIb emission of the 2FT-*o*CB dots along with the definite intrinsic noise in the detection system such as shot noise, read noise, etc. might be responsible for the evident decline of the mean SSIM index in NIR-IIb map. The highest consistency with the NIR-IIx+NIR-IIb detection in the NIR-II non-invasive visualization can be concluded and the advanced imaging technique is believed to hold the highest potential in the future clinical scene. As expected, the urine subsequently excreted from the mouse exhibits bright NIR-II emission, as shown in Supplementary Fig. 24f&g. After opening the abdomen, it could be seen that the bladder of the mouse was bulging with 2FT-*o*CB dots of mauve (see Supplementary Fig. 24h). Supplementary Fig. 25 gives a typical case of intraoperative bladder injury, where the perfused 2FT-*o*CB dots leak into the abdominal cavity and light up the whole body. Besides, the H&E staining results of the bladders in Supplementary Fig. 24i and Supplementary Fig. 26 show that there exists no obvious difference between the groups perfused with the heavy water dispersion of 2FT-*o*CB dots and the 1×PBS, respectively.

Computed tomography (CT) colonography has been widely utilized in clinics to examine the large intestine for cancer and growths called polyps by special x-ray equipment. The resulting ionizing radiation becomes the necessary cost to obtain an interior view of the colon which is otherwise only seen with a more invasive procedure by virtue of an endoscope. The fluorescence colonography was conducted with the assistance of 2FT-*o*CB dots, in different imaging windows of the NIR-II region after colonic perfusion (1 mg/mL, 200 μL, deuterium oxide dispersion). As shown in Supplementary Fig. 27a, integrated interaction between light and tissue in different imaging regions presents characteristic performance. Different degrees of diffused components blend in with the ballistic light, and they are both collected onto the near-infrared sensitive camera eventually. Some blurry edges and expanded outlines will undoubtedly interfere with the judgment and the operation. The NIR-IIx+NIR-IIb fluorescence image, by contrast, gives the most accurate visualization through the skin, muscle, fat, and other upper organs. To quantitatively divide the targeted area from the imaging background, the selected original results (white squares in Supplementary Fig. 27a) were processed by binarization, using three threshold values (TH = 0.27, 0.32 and 0.37, representing that 27%, 32%, and 37% of the highest brightness in the selected area were set as the threshold values for the signals/background determination). The overall trend is that the calculated area shrank with the red-shifting of the imaging wavelength, as shown in Supplementary Fig. 27b. Notably, the selected section of the colon is minimized in 1400-1700 nm. Meanwhile, the signal to background ratio (SBR) can be determined with the mean intensities of the segmented colon calculated as the signals, and the mean intensities of the rest regarded as the background. The measurement results of the three different THs were averaged and the data were presented as mean ± s.e.m. It can be patently seen in Supplementary Fig. 27b that the SBR value reaches the maximum with 1400-nm long-pass collection due to the background suppression. Eventually, the 2FT-*o*CB dots can be excreted from the mouse body, and the feces in the bright field image and the NIR-II fluorescence image under 793 nm laser excitation can be seen in Supplementary Fig. 27c&d. The bright field and NIR-II fluorescence images after opening the abdomen are shown in Supplementary Fig. 27e and Supplementary Fig. 28, and ensure the labeling of the colon. Meanwhile, the results of the histology study of the colons show no obvious side effects after the perfusion of 2FT-oCB dots in heavy water dispersion (see Supplementary Fig. 27f and Supplementary Fig. 29). The *in vivo* NIR-IIx+NIR-IIb fluorescence colonography is believed to possess tremendous potential for further clinical translation.

### Deep-penetration intrauterine residue detection

The uterine diseases significantly harmed female patients and a proper hysterography in the examinations of gynecology and obstetrics could guide to diagnose the intrauterine abnormity such as the endometrial polyp, adhesive uteritis, submucous myoma or adenomyosis, uterine deformity, etc. Fluorescence hysterography is free of ionizing radiation but is admittedly challenging since the uterus is always deeper than the bladder in the body and there usually exist adipose tissues surrounding uteruses and blocking the propagation of the optical signals, which raises the requirement of enough emission intensity for the nanoprobes. After uterine perfusion of the 2FT-*o*CB dots (1 mg/mL, 200 μL, deuterium oxide dispersion), the anesthetized mice laid on the imaging stage with the lower abdomen exposed to the irradiation of 793 nm CW laser. The NIR-II fluorescence uterus images are displayed in Supplementary Fig. 30a. The calculated diameter of the right uterus is minimized at 1.69 mm in the NIR-IIx+NIR-IIb image by the measurement of full width at half maxima (FWHM), which is demonstrated in Supplementary Fig. 30b&c, while the FWHM measured in NIR-IIb (1500-nm long-pass) image was 1.78 mm. It is worth mentioning that the uterus in the 900-nm long-pass image even shows higher spatial resolution (2.92 mm) compared with that (2.97 mm) in the 1000-nm long-pass image, which is consistent with our previous conclusion and again confirms the comparable imaging strength in 900-1000 nm owing to the light absorption peak at ∼980 nm. The visible brightness difference between the left and right can be blamed on the uneven distribution of the adipose and muscular tissues. After the enhancement on image brightness, it can be seen that the left uterus is also visualized and precisely positioned (see the inset images of Supplementary Fig. 30a). As shown in Supplementary Fig. 30d&e, the 2FT-*o*CB dots can pass out of the body with the constant peristalsis of the uteruses after imaging, and residual 2FT-*o*CB dots can also be removed by uterine lavage. Further demonstration about the labeling of uteruses can be seen in Supplementary Fig. 30f and Supplementary Fig. 31, and the bright 2FT-*o*CB dots from the uteruses exhibit strong NIR-II emission after opening the abdomen. Thus, the 2FT-*o*CB dots assisted *in vivo* fluorescence imaging with NIR-IIx+NIR-IIb detection is verified to provide precise spatial resolution with no invasion. Besides, the uterus treated with the heavy water dispersion of 2FT-oCB dots shows no noticeable damage in the H&E stained tissues (see Supplementary Fig. 30g and Supplementary Fig. 32).

After delivery, abortion, or surgery of suction and curettage, there might be some intrauterine residue remaining inside the uteruses, causing hemorrhage or infection. B-scan ultrasonography is widely used in the clinic to diagnose whether there is any residue, however, the small fragments remaining in the uterus are still difficult to identify. In parallel, NIR-II fluorescence imaging can give some new ideas for accurate detection with optical resolution. Within the spectral region around the absorption peak, the light absorption would deplete the scattered photons. Moreover, the residual tissues are always abundant in water, which leads to strong negative staining around the bright signals due to the light absorption. Thus, the dual contribution could provide a sharp contrast for further diagnosis. As shown in Fig. 4a, the super-absorbent resin is filled into the uterus cavities, acting as the tiny foreign bodies, such as hydatidiform moles. The *in vivo* imaging was conducted after abdominal suture and intrauterine injection of the 2FT-*o*CB dots (1 mg/mL, 200 μL, deuterium oxide dispersion) (see Fig. 4b), and the results can be seen in Fig. 4c. With the uterine peristalsis and auxiliary pressing, the tissue depth (about several millimeters, mainly consisting of the muscle and fat tissues), as well as the tissue interruption, reduces, and the intrauterine foreign bodies are increasingly identified. To better mimic the endogenous residue, a batch of pregnant mice were obtained (see Fig. 4d). After intrauterine injection of the 2FT-*o*CB dots (1 mg/mL, 500 μL, deuterium oxide dispersion), the fetuses in the uteruses with strong negative staining can be clearly presented (see Fig. 4e-g). The 1300LP, 1400LP, and 1500LP images were put into three channels with the pseudo-color enhancement of red, green, and blue, respectively. As shown in Fig. 4h, the merged image of the three channels shows generally white integrated by the three primary colors. However, an excess of purple integrated by the red and blue channels on the outer contours of some pregnancy tissues is also shown (especially in the yellow frame), which can be blamed on the relatively high light diffusion in the 1300LP and 1500LP channels. As shown in Fig. 4i, the placentas can be accurately outlined with NIR-IIx+NIR-IIb detection. Then, the detection of missed abortion was further performed. On 6^th^ day post injection of lipopolysaccharide (LPS, a kind of endotoxin, utilized to stimulate uterus to contract, and thus leading to abortion), the mice were imaged after intrauterine perfusion of the 2FT-*o*CB dots (1 mg/mL, 200 μL, deuterium oxide dispersion). From the NIR-IIx+NIR-IIb images *in vivo* shown in Fig. 4j, a fetus can be observed in the right uterus and some residual pregnancy tissue can be precisely detected in the left uterus. By positioning the boundary, the fetus is determined with a diameter of 5.45 mm (see Fig. 4k). Comparing the image before (see Fig. 4j&k) and after (see Fig. 4l) opening the abdomen, the hindrance of skin and adipose tissue does not disturb the precise diagnosis with NIR-IIx+NIR-IIb detection. As shown in Fig. 4m, the mouse with missed abortion after intrauterine perfusion of the 2FT-*o*CB dots looks no different from the normal mouse. Finally, the uteruses were further dissected out for section and H&E staining (see Fig. 4n). The section image of the right uterus shows a complete structure of the mouse fetus (see Fig. 4o and Supplementary Fig. 33), while the left side contains a mess of residual pregnancy tissue (see Fig. 4p and Supplementary Fig. 33), which is consistent with the *in vivo* NIR-IIx+NIR-IIb observation. The efficient labeling of fluorophores with bright NIR-IIx+NIR-IIb emission and the strong negative staining around the light absorption peak compose the dual visualization, which is believed to provide a powerful platform for the precise diagnosis of intrauterine residue.

**Figure 4.**
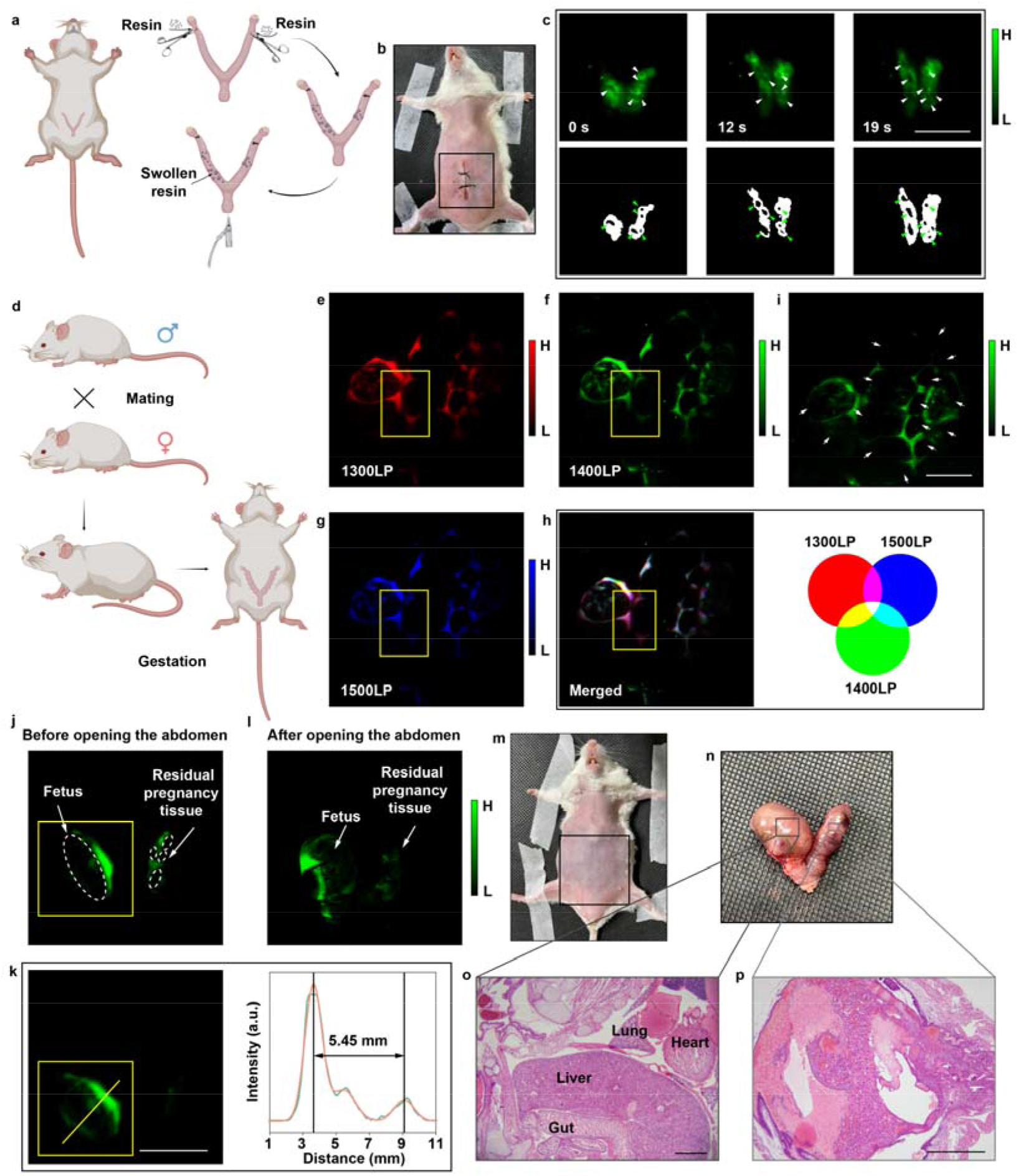
The precise detection of intrauterine foreign body *in vivo*. **(a)** The operation illustration for the intrauterine foreign body detection. **(b)** The picture of the mouse after operation with the abdomen sutured. **(c)** Top, the NIR-IIx+NIR-IIb fluorescence images with the peristalsis and manual pressing; Bottom, the binary images with the black spots representing the foreign bodies. The white arrows and the green arrows in (c) point out the foreign bodies. Scale bar, 10 mm. **(d)** The illustration of pregnancy. The fluorescence uterine images of pregnant mouse with **(e)** 1300-nm, **(f)** 1400-nm and **(g)** 1500-nm LP detection. **(h)** The merged image of red, green and blue channels. **(i)** The 1400LP image where the fetuses are marked with white arrows. Scale bar, 10 mm. **(j&k)** The fluorescence images of the missed abortion model before opening the abdomen. Scale bar, 10 mm. **(l)** The fluorescence imaging of the missed abortion model after opening the abdomen. The pictures of **(m)** the missed abortion mouse and **(n)** its isolated uteruses. Representative histological images of the **(o)** fetus and **(p)** residual pregnancy tissue. Scale bar in (o), 1 mm; Scale bar in (p), 500 μm.

## Conclusion

To ensure the absolute emission intensity beyond 1400 nm, a molecular design strategy of enhancing D-A interactions including reducing D-A distance and increasing electron donating ability based on the design of “backbone distortion and molecular rotors” was proposed, simultaneously ensuring both long absorption/emission wavelength and high fluorescence intensity in the aggregate state. The designed molecular dyes and dots show strong absorption properties and bright emission extending to 1700 nm. At the macroscopic level, the heavy water (deuterium oxide) dispersion can efficiently recover the fluorescence in the NIR-II region, especially beyond 1400 nm.

In the NIR fluorescence imaging, on the one side, the heavy water can efficiently improve the SBR. On the other side, the natural water (hydrogen oxide) in the animals could eliminate the scattering disturbance of biological tissue. Multiple evaluation methods are given in this work to fully confirmed the optimum fluorescence imaging quality with 1400-nm long-pass (NIR-IIx+NIR-IIb) detection. Taking full use of the absorption properties of water and heavy water, the non-invasive hollow organ fluorescence imaging can be performed with high quality. In addition, the lesions containing much water can be further negatively stained, thus making a stark contrast for precise diagnosis.

As a kind of secure isotopic composition, deuterium oxide exhibits no ionizing radiation and the dispersion with fluorophores shows almost no side effect on the mice. The intraluminal perfusion instead of intravenous injection shielded the organism from the introduced biosafety risk of the exogenous agents, which equipped the NIR-IIx+NIR-IIb fluorescence hollow organ imaging with tremendous potential for biological and clinical applications. Our future goal is to develop brighter fluorophores with remarkable emission near 1450 nm, completely withstanding the depletion in the NIR-IIx fluorescence imaging. We believe the pervasive optimization scheme for fluorescence imaging via molecular design, solution system improvement, and extending the imaging window can provide references for future researches.

## Methods

### Hydration and PEG-encapsulation of fluorescent molecules

A solution containing 3 mg of 2TT-oCB/2MTT-oCB/2MPT-oCB/2FT-oCB, 15 mg of Pluronic F-127, and 1.5 mL of tetrahydrofuran (THF) was added into 10 mL of deionized water. The mixture was then evenly stirred in the fume hood overnight to remove the THF thoroughly. The water dispersion of 2TT-oCB/2MTT-oCB/2MPT-oCB/2FT-oCB dots was then passed through a filter with the aperture of 0.22 μm for degerming and preliminary filtration. Then, after ultrafiltration, the dots were further purified and concentrated. The heavy water dispersion of dots was obtained by redispersion in heavy water after ultrafiltration.

### Animal handling

All experimental procedures were approved by Animal Use and Care Committee at Zhejiang University. The BALB/c nude mice (∼20 g) and Institute of Cancer Research (ICR) mice (∼20 g) involved in this work were provided from the SLAC Laboratory Animal Co. Ltd. (Shanghai, China). All the experimental animals were housed under standard conditions in the Laboratory Animal Center of Zhejiang University and had access to food and water ad libitum.

### NIR-II fluorescence macro imaging for the hollow organ *in vivo*

All the mice were anesthetized before the imaging via tribromoethanol (10 mL/kgBW of the 1.25% solution). After vesical, uterine, or colonic perfusion, the hollow organ fluorescence imaging was conducted with the corresponding organ exposed under the excitation of 793 nm CW laser (Suzhou Rugkuta Optoelectronics Co., Ltd., China). The dosages of 2FT-*o*CB dots (1 mg/mL) for the cystography, hysterography, and colonography were 20 μL, 200 μL, and 200 μL, respectively. The 900 nm, 1000 nm, 1100 nm, 1200 nm, 1300 nm, 1400 nm, and 1500 nm long-pass optical filters were purchased from Thorlabs. The simple schematic diagram of imaging system is shown in Supplementary Fig. 34. The NIR-II signals from the 2FT-*o*CB dots in the hollow organs are collected by the NIR antireflection fixed focus lens with a focal length of 35 mm (TKL35, Tekwin) through the optical filters and detected by the electronic-cooling 2D (640 pixels × 512 pixels) InGaAs camera (SD640, Tekwin). The power densities of the excitation and the exposure times in the *in vivo* experiments were listed in Supplementary Table 1.

### Image processing with the fast Fourier transform and inverse fast Fourier transformation

Two-dimensional fast Fourier transformation was utilized to acquire spatial frequency maps of selected areas in whole-body vessels imaging by MATLAB R2020b. The low-frequency components were placed in the four corner parts of the FFT image while the high-frequency components were shown near the center at first. To facilitate the observation, the low-frequency components were then shifted to the center and the high-frequency components were set to the four corners. The final image was processed by the natural logarithm. The whole image processing was shown in Supplementary Fig. 35.

The zero-frequency point in spatial frequency maps was set as the center of the circle and then statistical analysis was conducted by counting the values of all pixels in the specific radius, which represented the intensity of the specific spatial frequency in the original fluorescence images. A circular low-pass filter was designed to filter out the high-frequency components. The cut-off radius was adjusted to control cut-off frequency. Then inverse fast Fourier transformation was adopted to obtain the denoising images of the whole-body vessels.

### Image processing of the SSIM assessment

Some important perception-based facts including structural degradation, luminance masking as well as contrast masking collaborated in the SSIM. Taking the luminance (*l*), contrast (*c*), and the structure (*s*) into consideration, the expression for SSIM index could be described as:

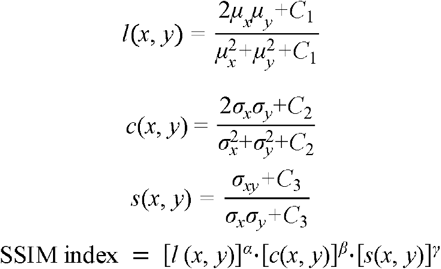

In the SSIM index, *l*(*x, y*) represents the comparison of the two images on brightness, *c*(*x, y*) differs the two images on contrast, and *s*(*x, y*) distinguishes the two images on the structural similarity and dissimilarity, where the *μ*_*x*_ and *μ*_*x*_ are the local means of intensity, *σ*_*x*_ and *σ*_*y*_ are the standard deviations and *σ*_*xy*_ is the cross-covariance for images *x* and *y* sequentially in the images. The constants *C*_1_ and *C*_2_ can be given directly from MATLAB to avoid the zero denominators, and *C*_2_ = 2*C*_3_. The presented NIR-II fluorescence bladder images were caught in their respective optimal shooting scene. The weight coefficient of brightness *α* was set to zero to exclude the systematic error caused by the experimental condition since the NIR-II fluorescence image brightness taken here contained the integrated influence of different exposure time and the discrepant emission intensity in the corresponding wavelength band under specific excitation. In comparison, the contrast and structure were not much impacted by the recording difference, so the weight coefficients of contrast (*β*) and structure (*γ*) were set to 1 empirically. Thus, the SSIM index in this work could be reduced as follows:

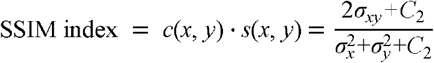

By invoking the built-in function in MATLAB R2020b, the SSIM map of the bladder image was generated, and the mean SSIM index of the whole map was calculated. SSIM map showed SSIM indexes of every pixel in the fluorescence image. In the SSIM assessment, the fluorescence bladder images through the skin, muscle, and adipose layer were set as the measured objects, and the bladder images after opening the abdomen were regarded as the references.

### Image processing of the binarization and segmentation

The selected areas in the fluorescence colon images were binarized with a given threshold value (TH) by invoking the built-in function in MATLAB R2020b. The threshold value, representing the proportion of the highest brightness in the selected area, determined the boundary of the signals and background, which excluded the noise disturbance to a certain extent. The areas of colon signals were calculated by counting the number of pixels in the binarized signal regions. The mean intensity of the binarized signals region was divided by the mean intensity of the binarized background region to calculate the SBR.

### Image processing of the FWHM analysis

The cross-sectional fluorescence intensity profiles along the lines over the uteruses were acquired by Image J. Nonlinear curve fitting of the intensity profiles was executed by the Gauss fitting in OriginPro 2018C, giving the calculated FWHM. In the SBR calculation, the peak of the fitted Gauss curve was recognized as the signal value, while the baseline of the fitted curve was considered as the background value.

### Preparation of the animal model for intrauterine foreign body detection

Female mice were anesthetized and the abdominal hair was removed. First, each mouse was placed in the supine position for laparotomy. Next, the right uterus was isolated and an incision was made in the right uterine horn. The uterine cavity was then filled with super absorbent resin through the incision. After that, the uterus and abdomen were sutured and 200 μL PBS was injected into the uterine cavity through the cervix to swell the resin. About 30 minutes later, the deuterium oxide dispersion of 2FT-*o*CB dots (1 mg/mL, 200 μL) was perfused into the uterus for fluorescence hysterography.

Another group of females were injected intraperitoneally with 10 IU/mouse of pregnant mare’s serum gonadotropin (PMSG) and received 10 IU/mouse intraperitoneal injection of human chorionic gonadotropin (HCG) 48 hours later. Then they were caged with males (two females per male). The vaginal plug was checked the following morning to indicate the day 0.5 of gestation (GD 0.5). Pregnant females at GD 12.5 were anesthetized and received 500 μL deuterium oxide dispersion of 2FT-*o*CB dots (1 mg/mL) via intrauterine perfusion by 26 G I.V. catheter (Jiangxi Fenglin, China) after abdominal hair removal for the next *in vivo* fluorescence imaging.

0.5 mg/kgBW LPS (Sigma-Aldrich, USA) was injected into the uteruses of the pregnant mice by intrauterine perfusion at GD 12.5 to induce abortion under anesthesia. The survival fetus and residual tissue were then examined by fluorescence hysterography after intrauterine perfusion of 2FT-*o*CB dots (1 mg/mL, 200 μL, deuterium oxide dispersion) at GD 18.5.

## Acknowledgment

This work was supported by the National Natural Science Foundation of China (61975172 and 82001874), Fundamental Research Funds for the Central Universities (2020-KYY-511108-0007), and Natural Science Foundation of Zhejiang Province (LR17F050001). The authors thank Chenyu Yang in the Center of Cryo-Electron Microscopy (CCEM), Zhejiang University for her technical assistance on Transmission Electron Microscopy.

