## supplementary figures and table for "Fluorescence visualization of deep-buried hollow organs"

**Materials.** All the chemicals and reagents were purchased from chemical sources, and the solvents for chemical reactions were distilled before use. All air and moisture sensitive reactions were carried out in flame-dried glassware under a nitrogen atmosphere.

**Measurements.** The UV-Vis-NIR absorption spectra measurement were performed using a Shimadzu UV-3600 spectrophotometer. $^{1}H$ and $^{13}C$ spectra were recorded at room temperature on a Unity-400 NMR spectrometer using $CDCl_3$ as solvent and tetramethylsilane (TMS) as a reference. Mass spectra (MS) were measured with a GCT premier CAB048 mass spectrometer in MALDI-TOF mode. The photoluminescence (PL) spectra were measured via an Ideaoptics NIR2200 spectrofluorometer. Dynamic light scattering (DLS) was measured on a Zetasizer Nano-ZS analyzer. Transmission electron microscopy (TEM) images were acquired from a Tecnai™ Spirit transmission electron microscope with an accelerating voltage of 120 kV. Density functional theory (DFT) calculations were carried out by the B3LYP/6G(d), Gaussian 09 package.

**Synthetic procedures and characterization data for the compounds.**

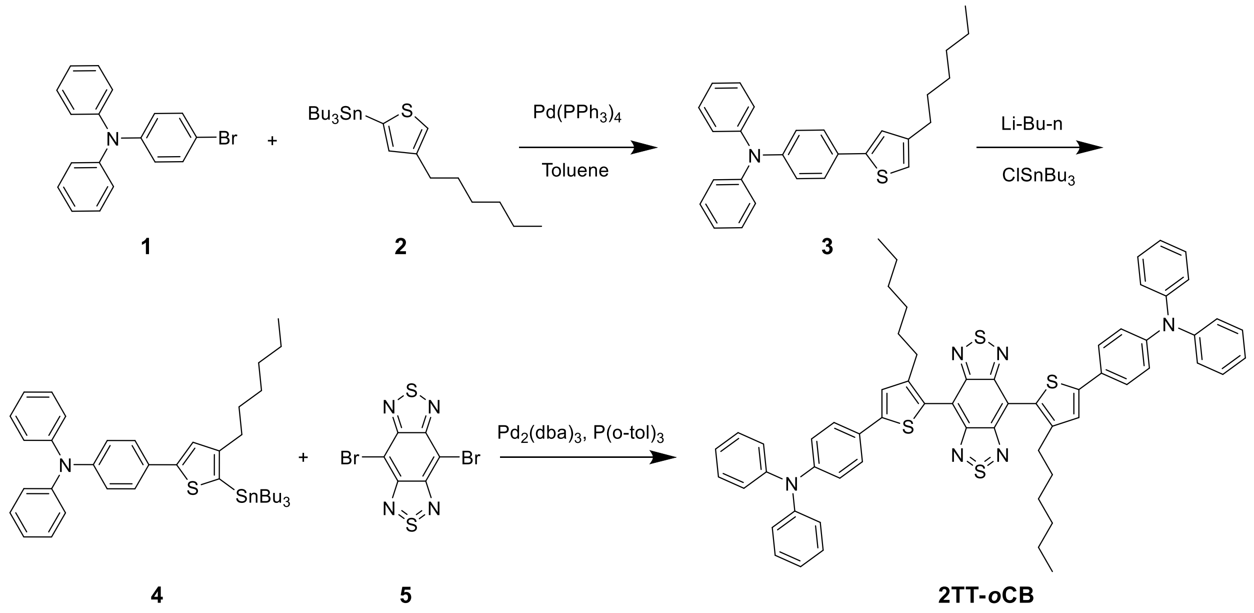


**Supplementary Scheme 1.** Synthetic route of **2TT-*o*CB**.

**Synthetic route of 3.**

Under $N_2$ atmosphere, **1** (1 g, 3.1 mmol), **2** (1.42 g, 3.1 mmol), $Pd(PPh_3)_4$ (180 mg, 0.15 mmol), and 20 mL toluene were added to a 100 mL predried two-necked flask. The mixture was refluxed for 24 h. After cooling down to room temperature, the solvent was removed by rotary evaporation. The crude product was purified by silica gel column to obtain the target molecule (yield, 70%). $^{1}H$ NMR (400 MHz, $CDCl_3$) δ 7.45 (2H, d, J = 8.8 Hz), 7.28-7.24 (4H, 8m), 7.12-7.10 (4H, m), 7.07-7.01 (5H, m), 2.60 (2H, t, J = 7.7 Hz), 1.67-1.60 (2H, m), 1.38- 1.27 (6H, m), 0.89 (3H, t, J = 6.7 Hz).

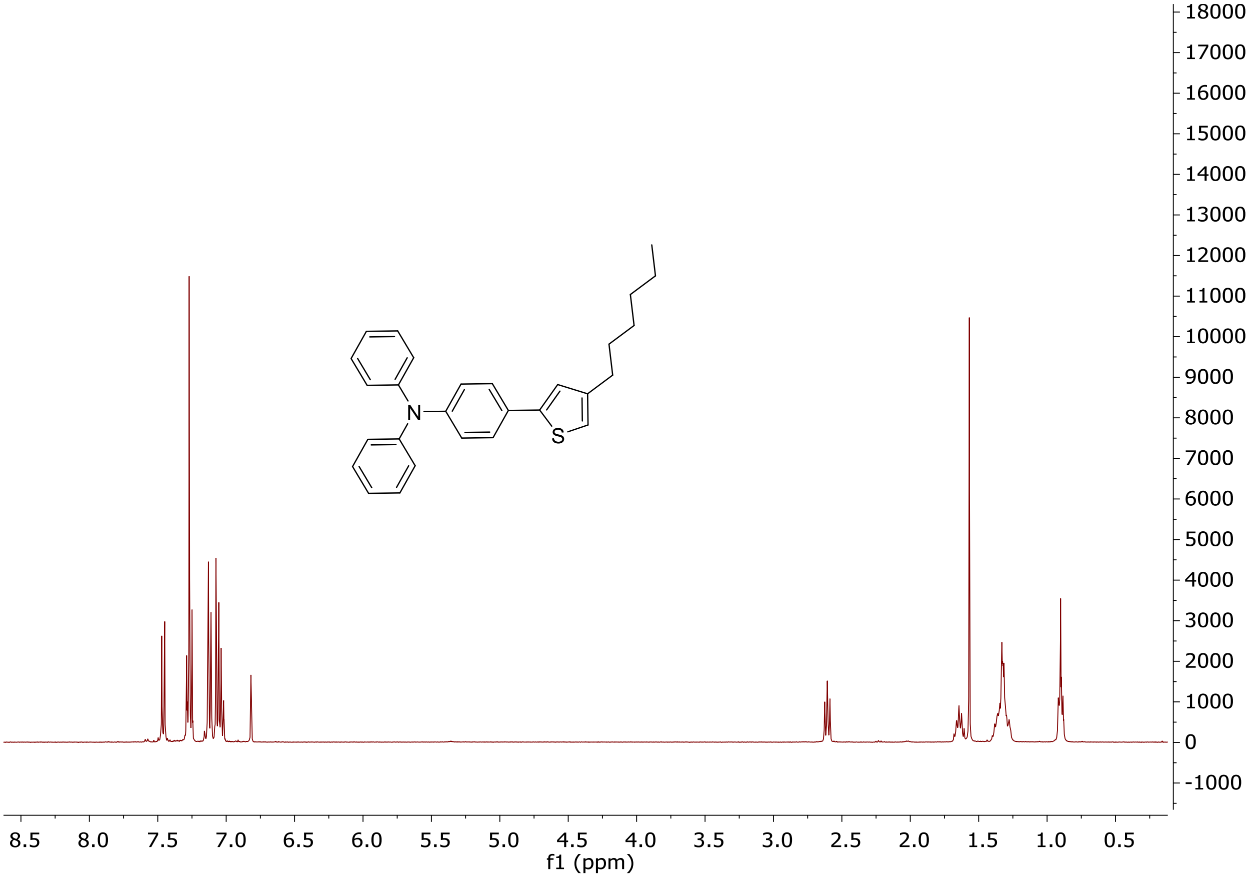


**Supplementary Fig. 1.** $^1$H NMR spectrum of **3**.

**Synthetic route of 4.**

nBuLi (2.3 mL, 5.8 mmol, 2.4 M in hexane) was added dropwise to a solution of **3** (2.16 g, 5.2 mmol) in THF (30 mL) at -78 °C. The reaction mixture was stirred 1 h at -78 °C. Then tributyltin chloride (1.8 g, 5.8 mmol) was added into the reaction at one portion. After stirring the mixture for 12 h at room temperature, KF solution was added to quench the reaction. The mixture was extracted with hexane three times, the combined organic phase was dried with $Na_2SO_4$. After removing the solvent, the product was used directly without further purification.

**Synthetic route of 2TT-*o*CB.**

A two-necked flask was charged with **4** (0.7 g, 1 mmol), **5** (87 mg, 0.25 mmol), $Pd_2(dba)_3$ (22 mg, 0.025 mmol), P(*o*-tol)$_3$ (66 mg, 0.21 mmol), and degassed dry toluene (1.5 mL), and heated overnight under 110 °C. Upon cooling, the crude product was quenched with KF solution and extracted with DCM. The combined organic phase was dried with $Na_2SO_4$. After removing the solvent, the product was purified with silica column to obtain a dark green solid (yield: 45%). $^1$H NMR (400 MHz, $CDCl_3$), δ (ppm) = 7.59-7.56 (4H, m), 7.37 (2H, s), 7.31-7.26 (8H, m), 7.16-7.12 (8H, m), 7.11-7.03 (8H, m), 2.61-2.57 (4H, t, J = 8Hz), 1.63, (4H, m), 1.15-1.10 (12H, m), 0.73 (6H, m). $^{13}$C NMR (100 MHz, $CDCl_3$), δ (ppm): 152.6, 146.9, 146.8, 146.3, 145.0, 128.7, 127.5, 126.1, 124.1, 124.0, 122.7, 122.5, 115.4, 99.3, 30.8, 29.8, 29.6, 28.4, 21.8, 13.3. MS: m/z: [M]+ calcd for $C_{62}H_{56}N_6S_4$: 1012.3, found: 1012.3.

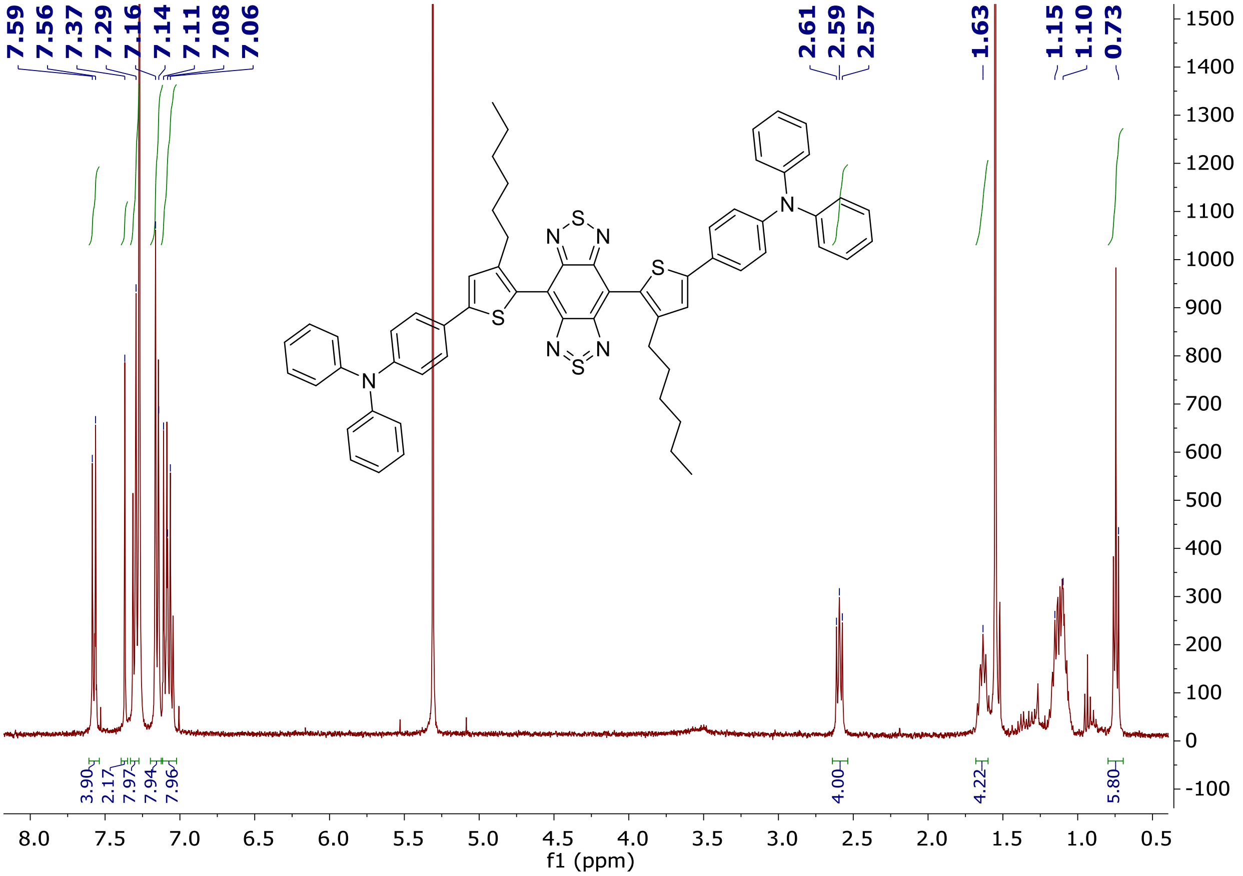


**Supplementary Fig. 2.** $^{1}$H NMR spectrum of **2TT-*o*CB**.

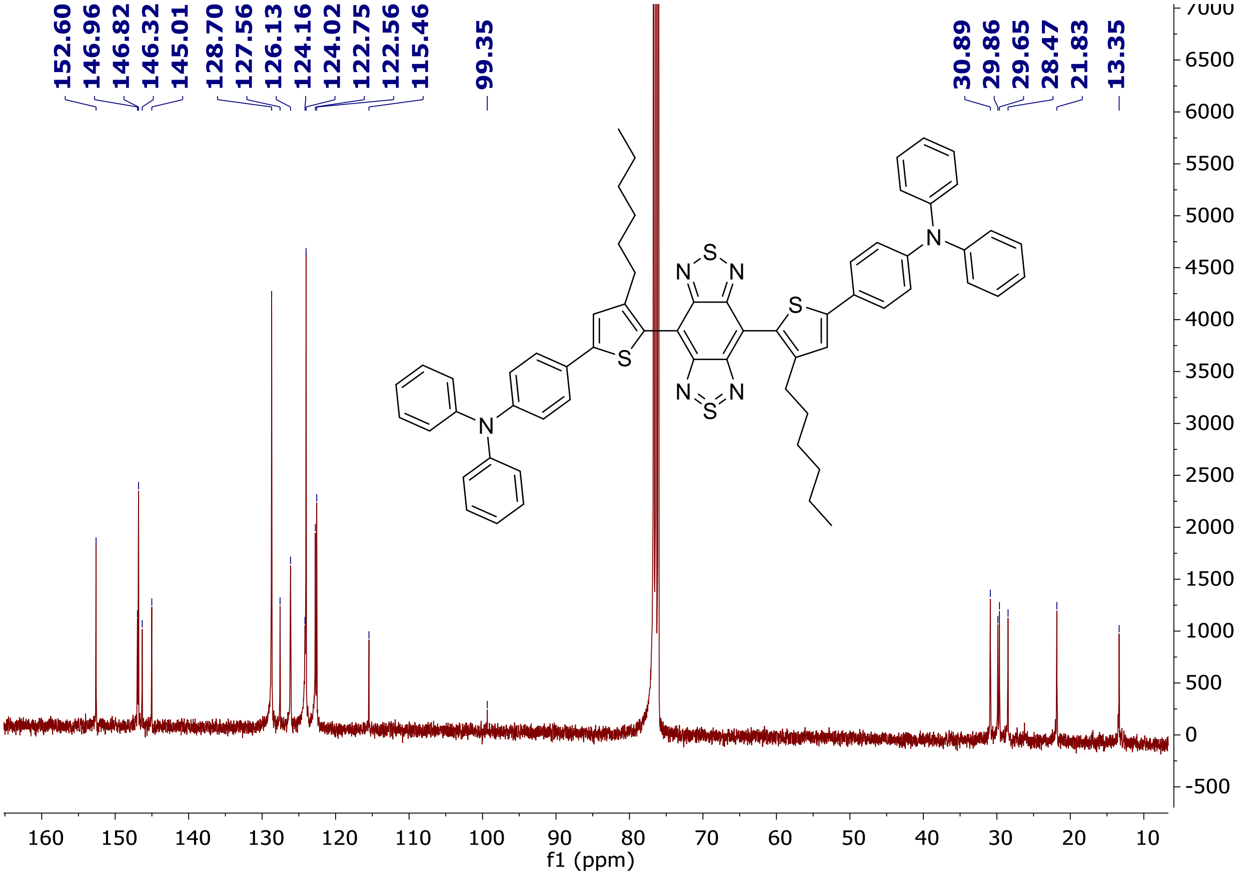


**Supplementary Fig. 3.** $^{13}$C NMR spectrum of **2TT-*o*CB**.

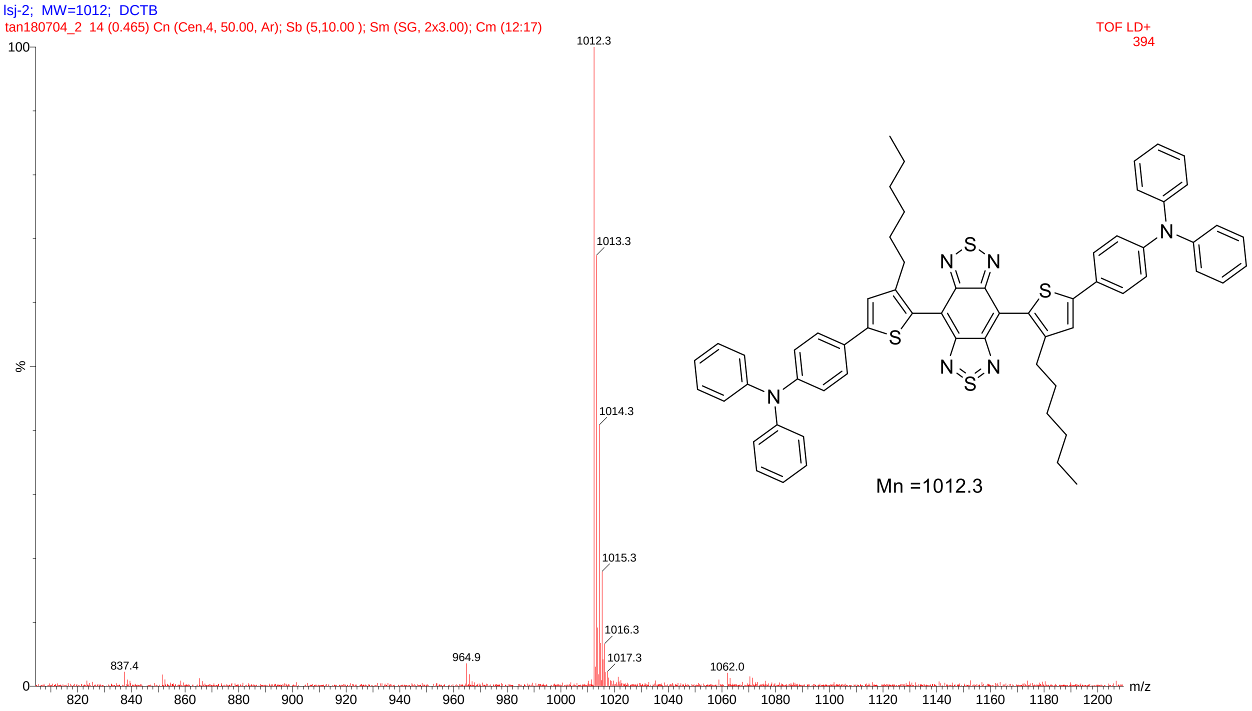


**Supplementary Fig. 4.** MALDI-TOF-MS spectrum of **2TT-*o*CB**.

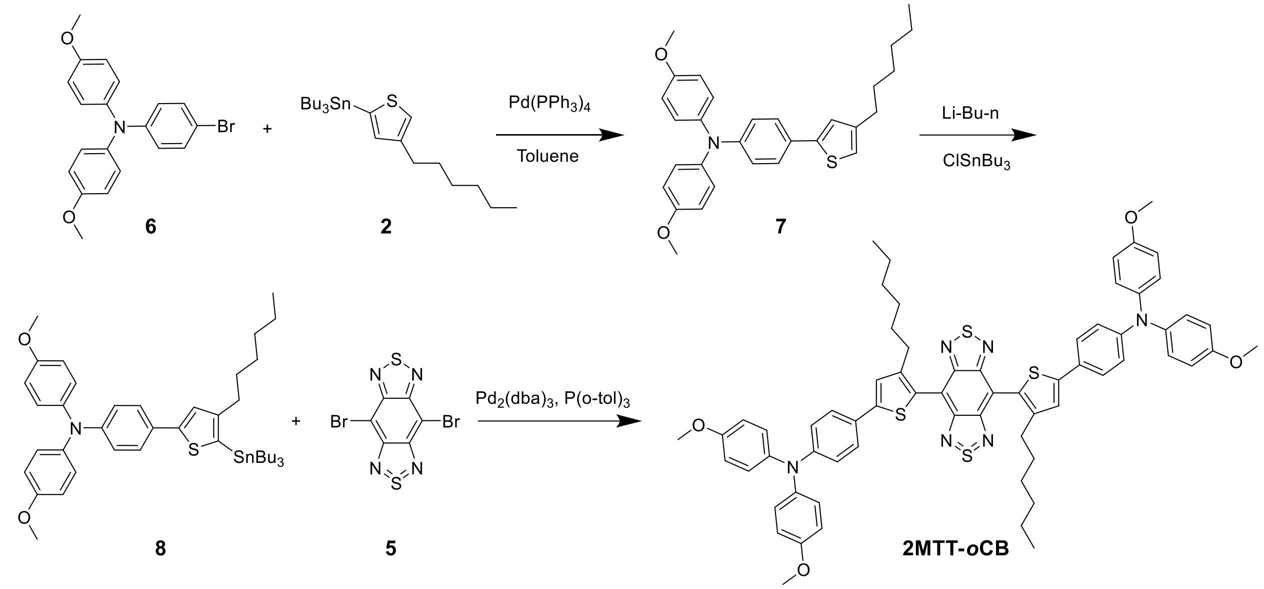


**Supplementary Scheme 2.** Synthetic route of **2MTT-*o*CB**.

**Synthetic route of 2MTT-*o*CB.**

The synthetic route to MTT1-*o*CB was similar to that of TT1-*o*CB by changing **1** into **6**. $^{1}$H NMR (400 MHz, $CDCl_3$), δ (ppm) = 7.54 (4H, m), 7.37 (2H, s), 7.12 (8H, m), 6.96 (12H, m), 6.88 (4H, m), 3.84 (12H, s), 2.62 (4H, m), 1.67 (4H, m), 1.12 (12H, m), 0.77 (6H, m). $^{13}$C NMR (100 MHz, $CDCl_3$), δ (ppm): 156.05, 153.23, 148.53, 147.37, 145.62, 140.60, 127.76, 126.63, 126.27, 124.34, 120.23, 116.06, 114.77, 55.51, 31.54, 30.53, 30.30, 29.13, 22.49, 14.03.

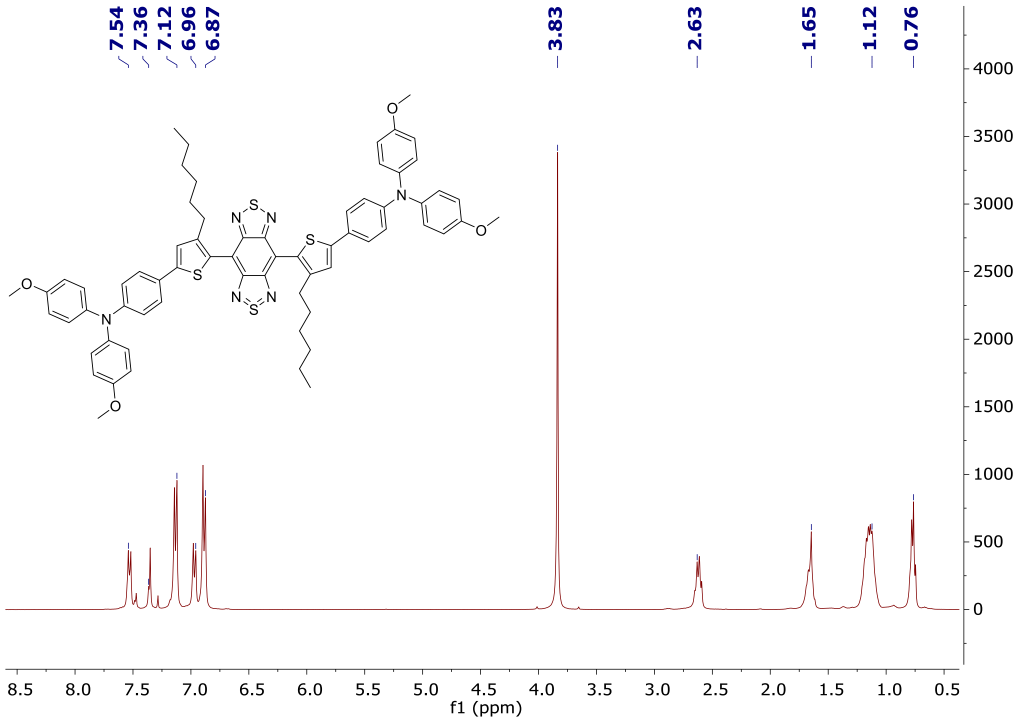


**Supplementary Fig. 5.** $^{1}$H NMR spectrum of **2MTT-*o*CB**.

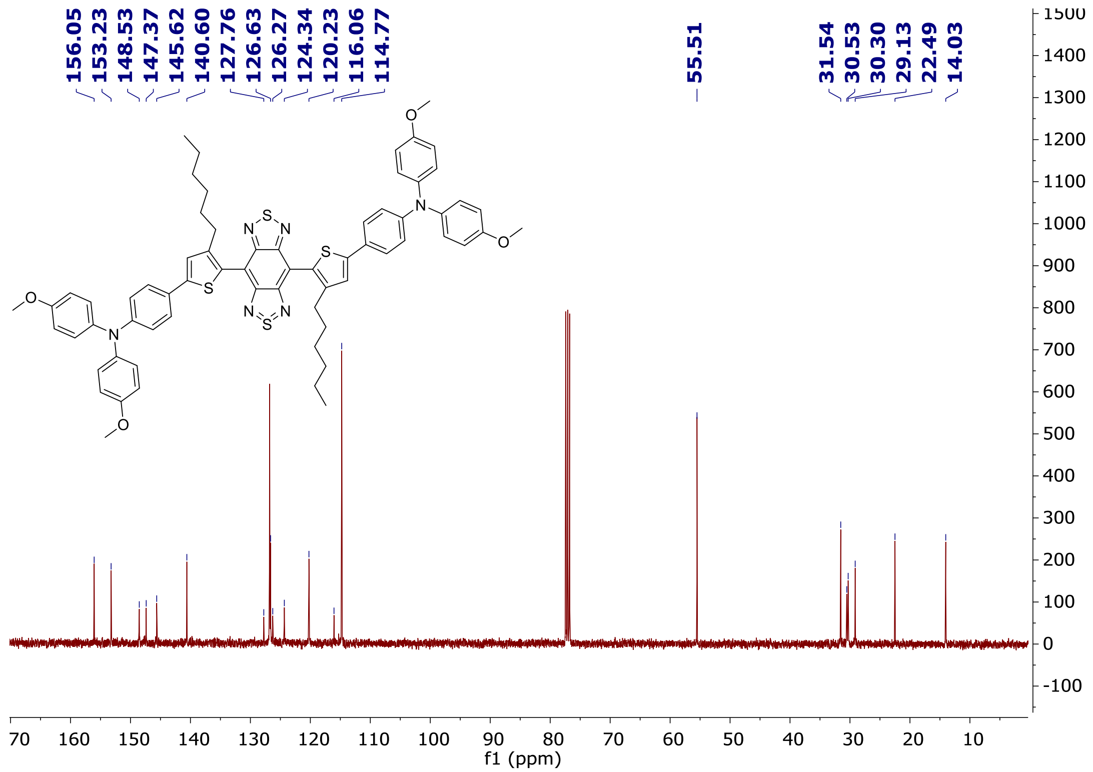


**Supplementary Fig. 6.** $^{13}$C NMR spectrum of **2MTT-*o*CB**.

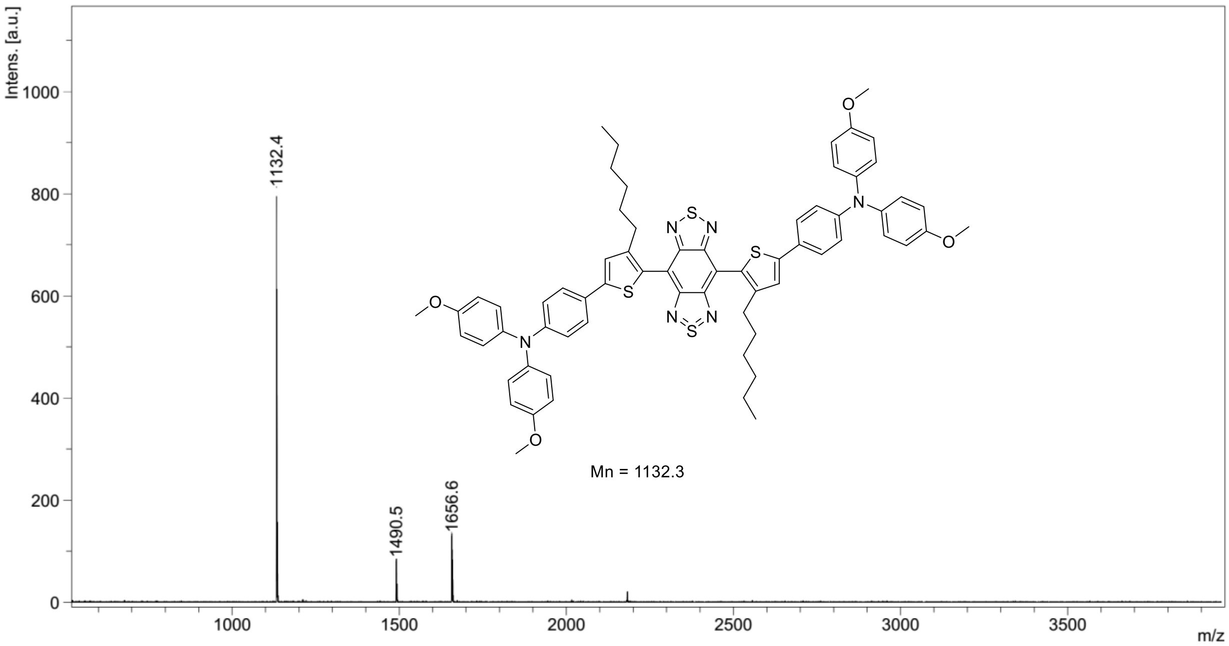


**Supplementary Fig. 7.** MALDI-TOF-MS spectrum of **2MTT-*o*CB**.

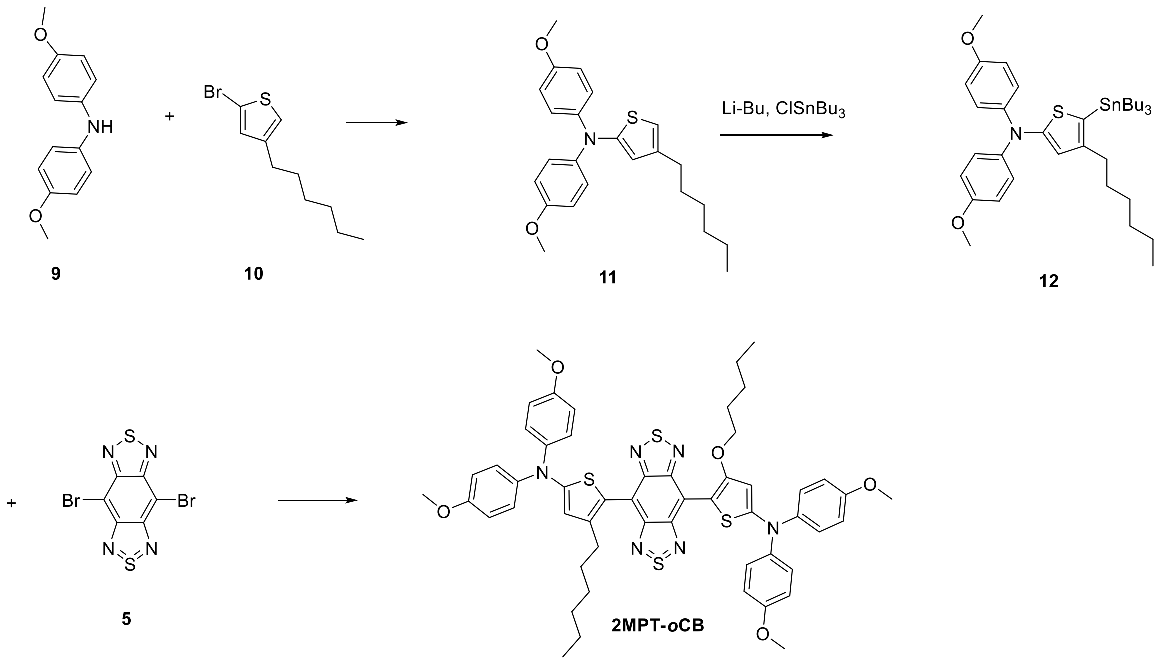


**Supplementary Scheme 3.** Synthetic route of **2MPT-*o*CB**.

**Synthetic route of 11.**

Bis(4-methoxyphenyl)amine (**9**, 0.23 g, 1 mmol), 2-bromo-4-hexylthiophene (**10**, 0.25 g, 1 mmol), $Pd_2(dba)_3$ (46 mg, 0.05 mmol), $P(t\text{-}Bu)_3$ (0.8 mL, 0.4 mmol, 10w/v% in pentane), NaOBu-*t* (21.3 mg, 1.3 mmol) and toluene (10 mL) were added into a two-necked flask. The mixture was refluxed for 24 h under protection of nitrogen. After cooling down to room temperature, water was added to quench the reaction and the organic phase was extracted and dried. The crude product was purified by a silica gel column to obtain the product (yield: 70%). $^1H$ NMR (400 MHz, $CDCl_3$), δ (ppm) = 7.11 (4 H, m), 6.85 (4H, m), 6.45 (1H, s), 6.42 (1H, s), 3.83 (6H, s), 2.50 (2H, t, J = 8 Hz), 1.62 (2H, m), 1.36 (6H, m), 0.92 (3H, m).

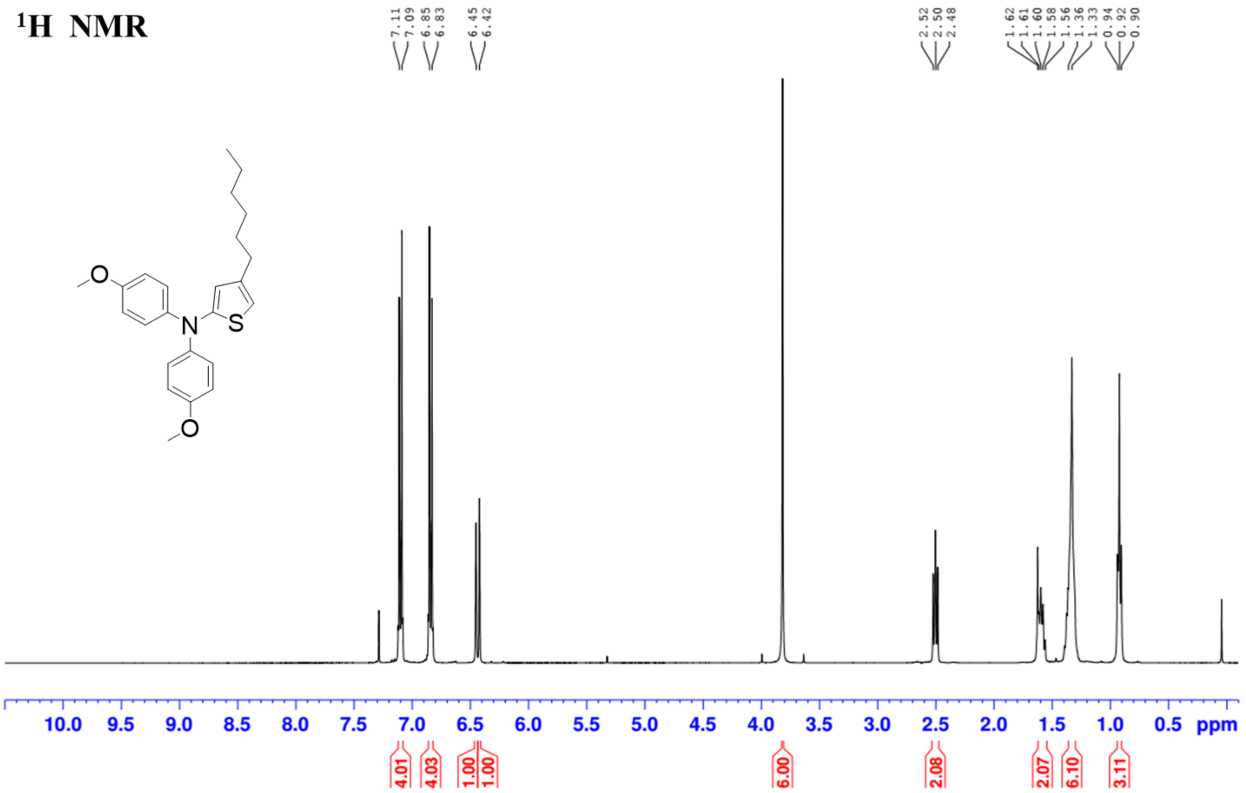


**Supplementary Fig. 8.** $^{1}$H NMR spectrum of **11**.

**Synthetic route of 12.**

nBuLi (2.3 mL, 5.8 mmol, 2.4 M in hexane) was added dropwise to a solution of **11** (2.05 g, 5.2 mmol) in THF (30 mL) at -78 °C. The reaction mixture was stirred 1 h at -78 °C. Then tributyltin chloride (1.8 g, 5.8 mmol) was added into the reaction at one portion. After stirring the mixture for 12 h at room temperature, KF solution was added to quench the reaction. The mixture was extracted with hexane three times, the combined organic phase was dried with $Na_2SO_4$. After removing the solvent, the product was used directly without further purification.

**Synthetic route of 2MPT-*o*CB**.

The synthetic route to 2MPT-*o*CB was similar to that of 2MTT-*o*CB by changing **8** into **12**. $^{1}$H NMR (400 MHz, $CDCl_3$), δ (ppm) = 7.32 (8H, d, J = 8Hz), 6.90 (8H, d, J = 8Hz), 6.49 (2H, m), 3.82 (12H, s), 2.50 (4H, t, J = 8Hz), 1.52 (4H, m), 1.12 (12H, m), 0.76 (6H, m). $^{13}$C NMR (100 MHz, $CDCl_3$), δ (ppm): 157.37, 156.56, 153.20, 144.29, 140.74, 126.14, 115.14, 114.72, 114.64, 55.50, 31.54, 31.05, 29.21, 22.52, 14.02. MS: m/z: [M]+ calcd for $C_{54}H_{56}N_6S_4O_4$: 980.3, found: 980.3.

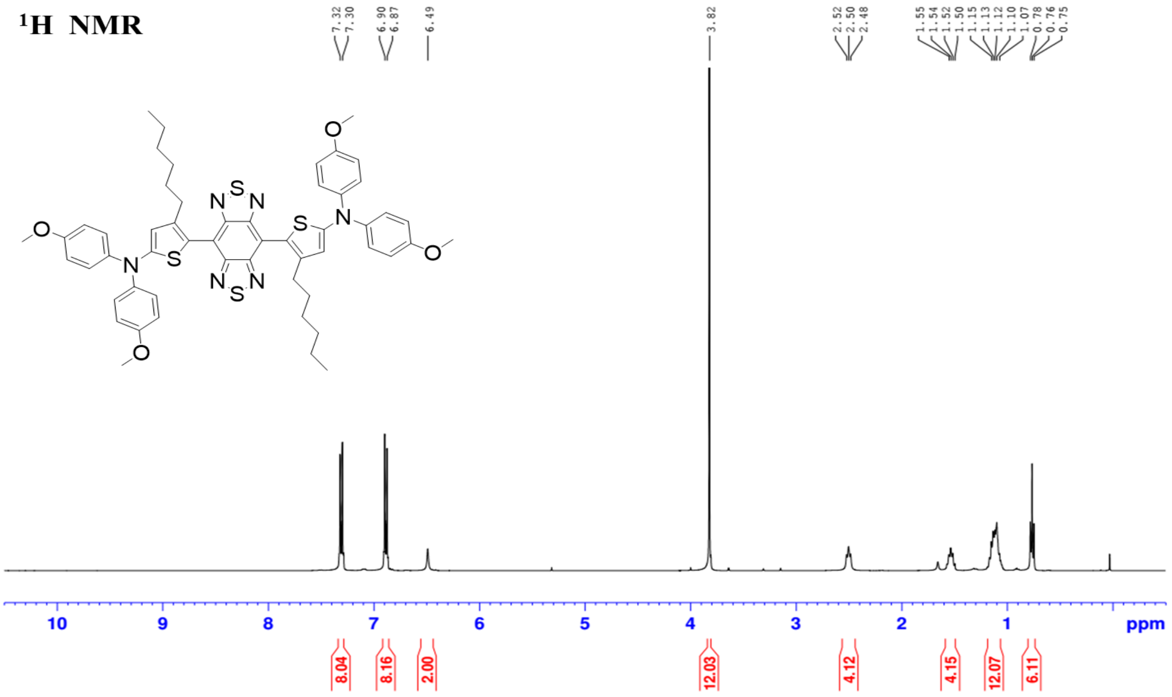


**Supplementary Fig. 9**. $^{1}$H NMR spectrum of **2MPT-*o*CB**.

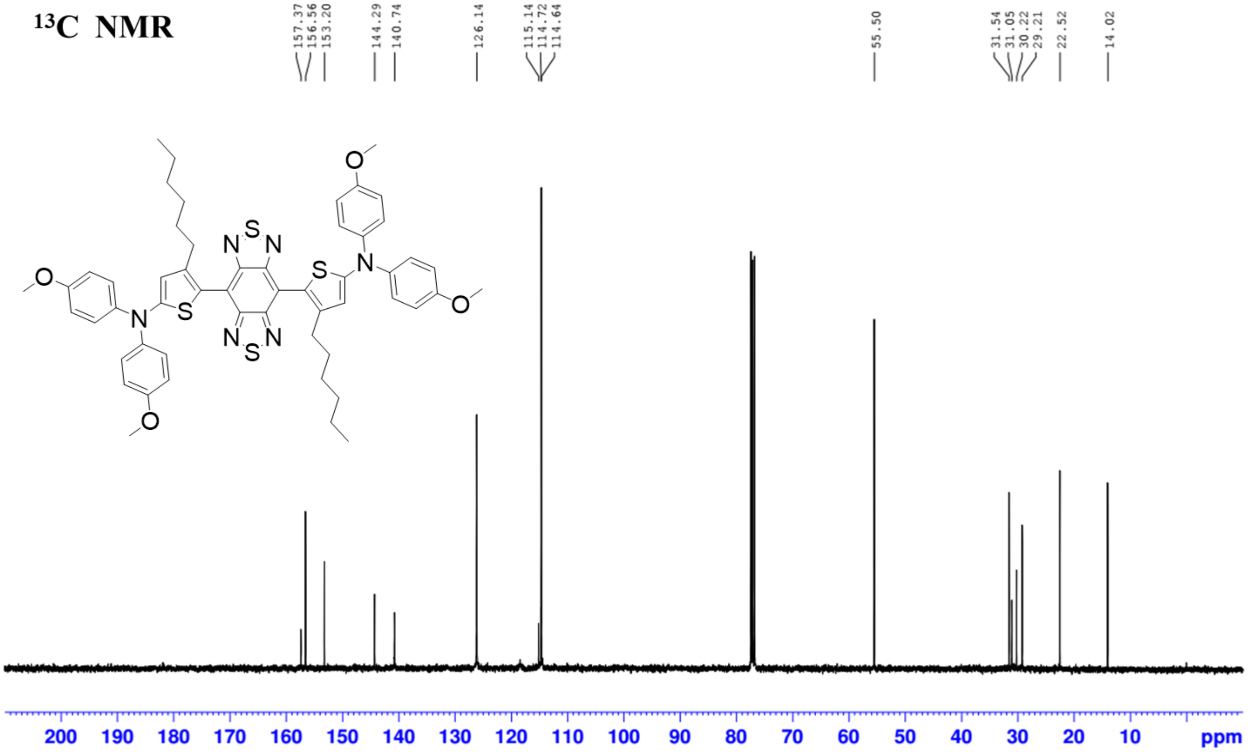


**Supplementary Fig. 10**. $^{13}$C NMR spectrum of **2MPT-*o*CB**.

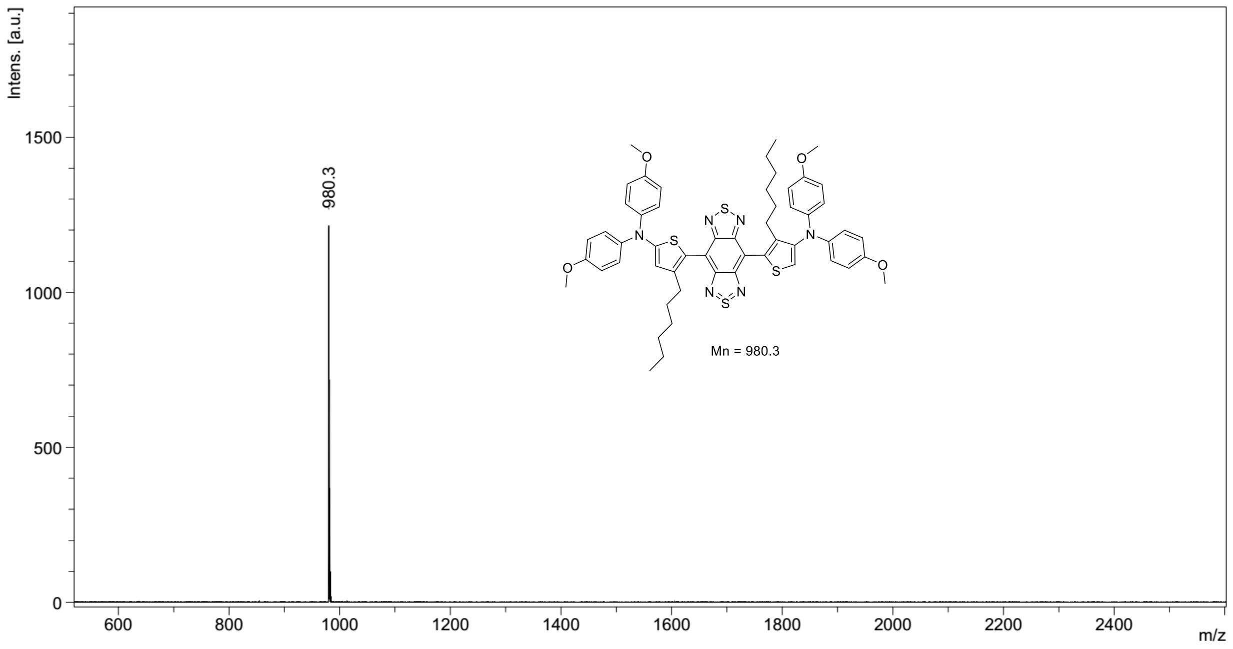


**Supplementary Fig. 11**. MALDI-TOF-MS spectrum of **2MPT-*o*CB**.

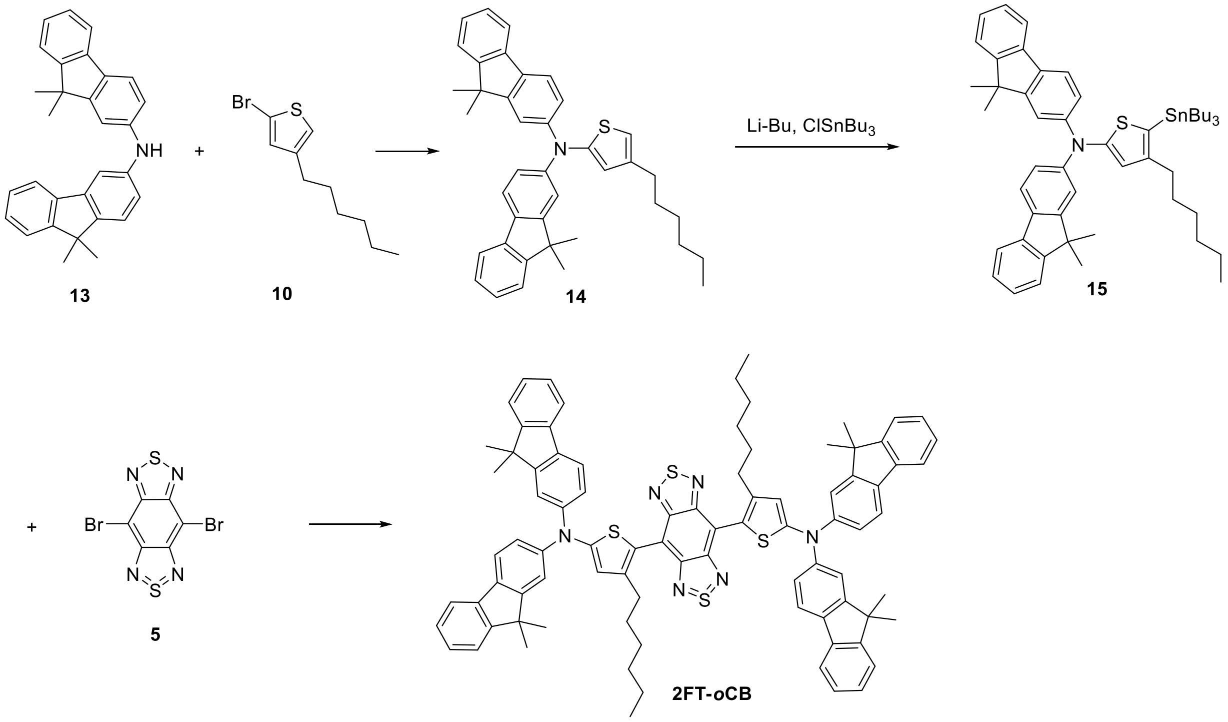


**Supplementary Scheme 4**. Synthetic route of **2FT-*o*CB**.

**Synthetic route of 14.**

2-bromo-4-hexylthiophene (**10**, 0.25 g, 1 mmol), **13** (0.56 g, 1 mmol), $Pd_2(dba)_3$ (46 mg, 0.05 mmol), P(t-Bu)$_3$ (0.8 mL, 0.4 mmol, 10w/v% in pentane), NaOBu-*t* (21.3 mg, 1.3 mmol) and toluene (10 mL) were added into a two-necked flask. The mixture was refluxed for 24 h under protection of nitrogen. After cooling down to room temperature, water was added to quench the reaction and the organic phase was extracted and dried. The crude product was purified by a silica gel column to obtain the product (yield: 71%). $^1$H NMR (400 MHz, $CDCl_3$), δ (ppm) = 7.68 (8H, m), 7.44 (2H, m), 7.32 (2H, m), 7.16 (2H, m), 6.68 (1H, s), 6.61 (1H, s), 2.58 (2H, t, J = 8Hz), 1.64 (2H, m), 1.46 (12H, s), 1.35 (6H, m), 0.93 (3H, m).

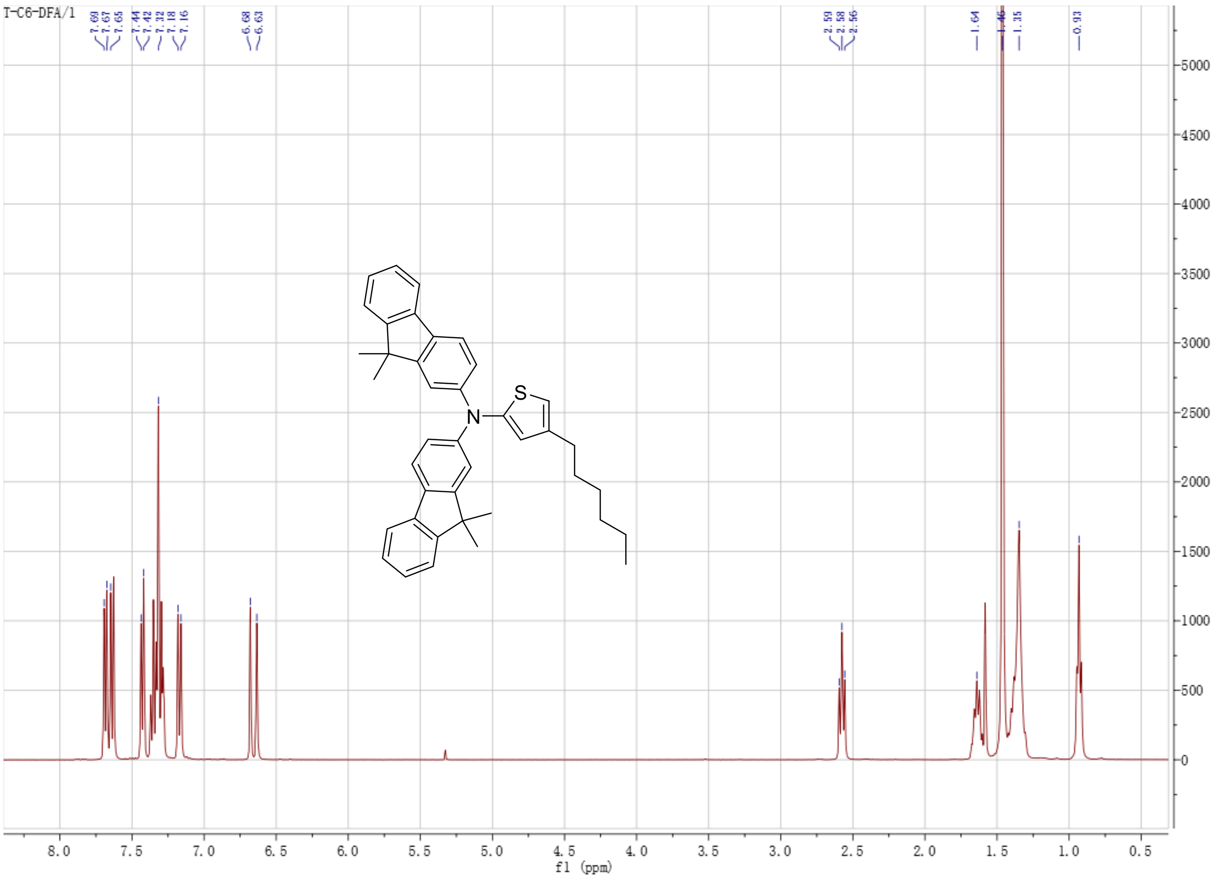


**Supplementary Fig. 12**. $^1$H NMR spectrum of **14**.

**Synthetic route of 15.**

nBuLi (2.3 mL, 5.8 mmol, 2.4 M in hexane) was added dropwise to a solution of **14** (2.94 g, 5.2 mmol) in THF (30 mL) at -78 °C. The reaction mixture was stirred 1 h at -78 °C. Then tributyltin chloride (1.8 g, 5.8 mmol) was added into the reaction at one portion. After stirring the mixture for 12 h at room temperature, KF solution was added to quench the reaction. The mixture was extracted with hexane three times, the combined organic phase was dried with $Na_2SO_4$. After removing the solvent, the product was used directly without further purification.

**Synthetic route of 2FT-*o*CB.**

The synthetic route to **2FT-*o*CB** was similar to that of **2MPT-*o*CB** by changing **12** into **15**. $^1$H NMR (400 MHz, $CDCl_3$), δ (ppm) = 7.77 (4H, m), 7.54 (2H, s), 7.47-7.28 (22H, m), 2.63 (4H, t, J = 8Hz), 1.63 (4H, m), 1.52 (24H, m), 1.21 (12H, m), 0.80 (6H, m). $^{13}$C NMR (100 MHz, $CDCl_3$), δ (ppm): 155.60, 155.01, 153.69, 153.28, 146.99, 144.07, 138.84, 135.32, 127.09, 126.80, 123.18, 122.58, 120.71, 119.66, 118.60, 115.37, 68.02, 46.99, 31.60, 31.12, 30.40, 29.20, 27.11, 25.68, 22.54, 14.07. MS: m/z: [M]+ calcd for $C_{86}H_{80}N_6S_4$: 1324.5, found: 1324.5.

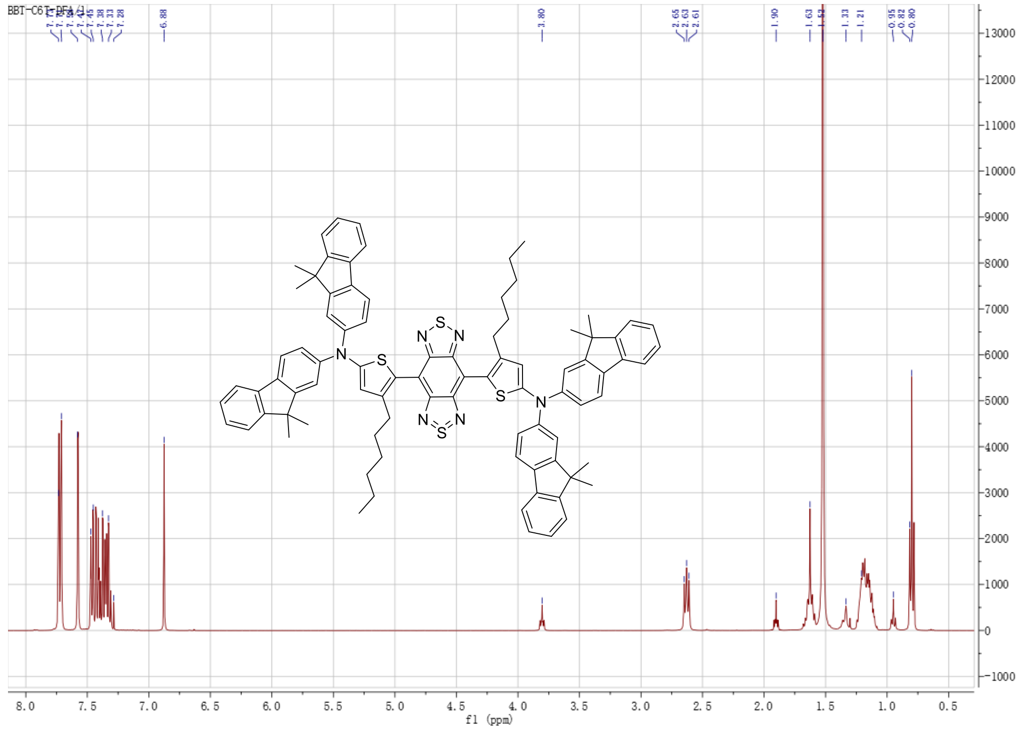

**Supplementary Fig. 13**. $^{1}$H NMR spectrum of **2FT-*o*CB**.

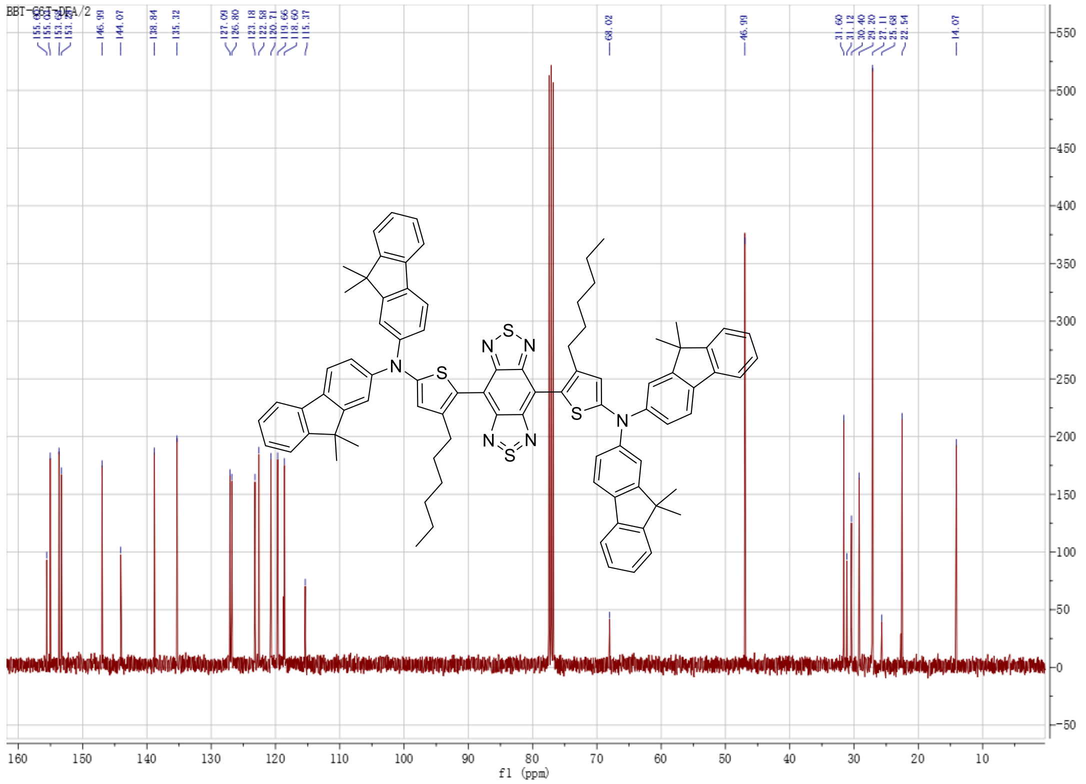

**Supplementary Fig. 14**. $^{13}$C NMR spectrum of **2FT-*o*CB**.

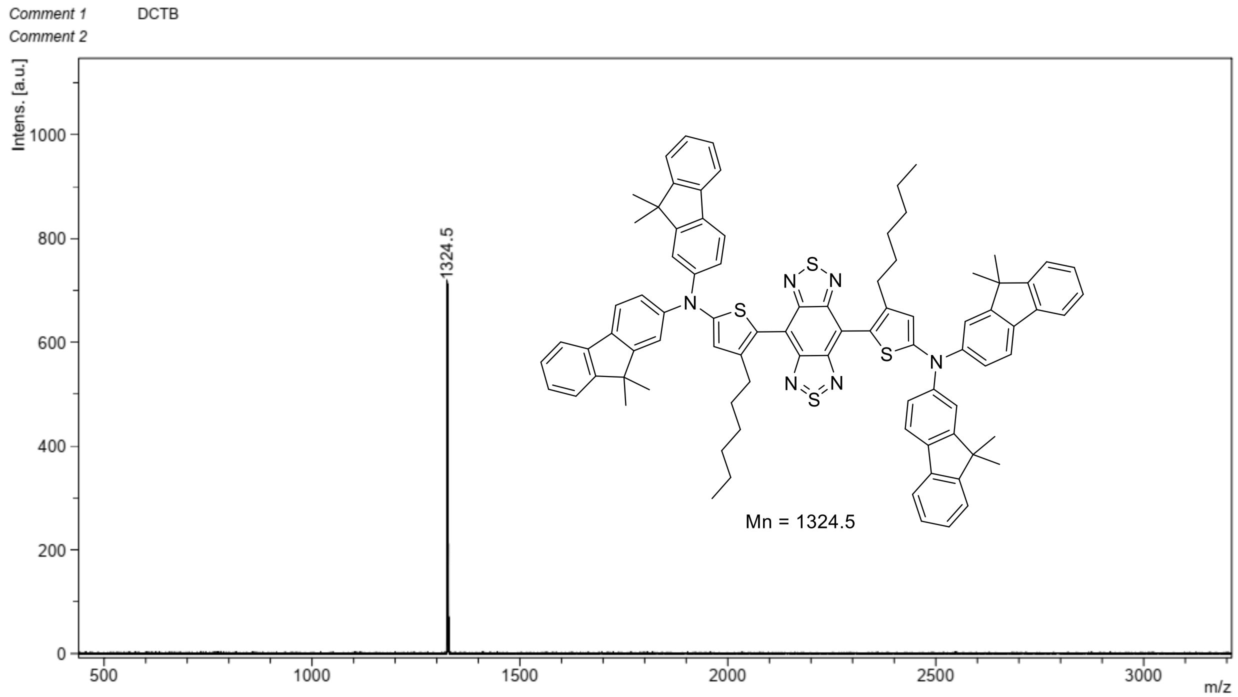


**Supplementary Fig. 15**. MALDI-TOF-MS spectrum of **2FT-*o*CB**.

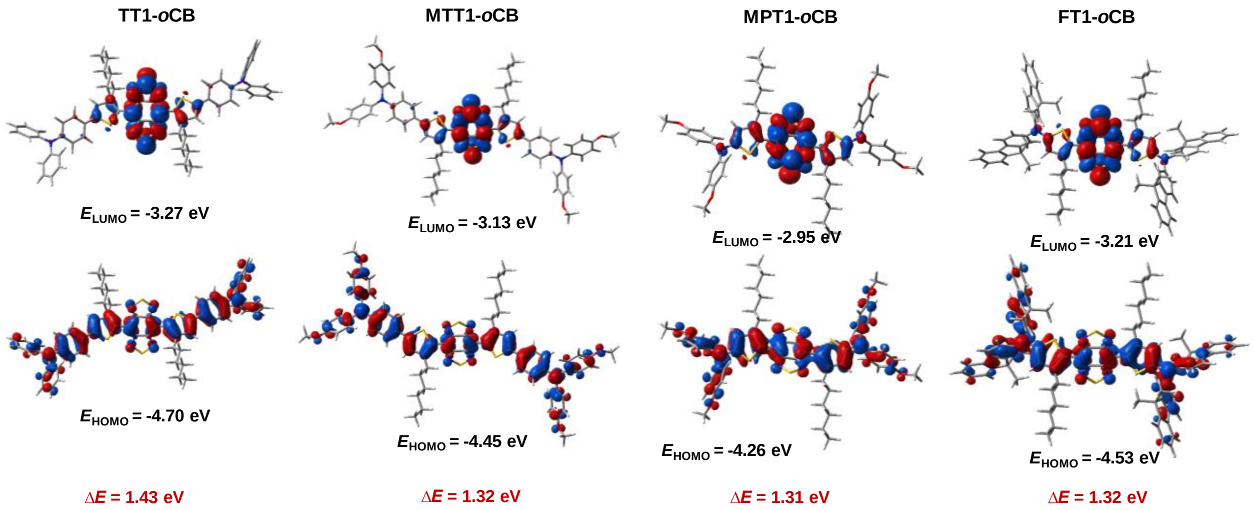


**Supplementary Fig. 16**. The frontier molecular orbitals of the lowest unoccupied molecular orbital (LUMO) and the highest occupied molecular orbital (HOMO) determined at the B3LYP/6-311g(d,p) level of theory.

The HOMO of the molecules is delocalized along the whole molecule, revealing an excellent molecular conjugation. While the LUMO is primarily located on the BBTD core. The energy gap between HOMO and LUMO for 2TT-*o*CB (1.43 eV), 2MTT-*o*CB (1.32 eV), 2MPT-*o*CB (1.31 eV) and 2FT-*o*CB (1.32 eV) decreases with an increase of D-A interactions, indicating enhanced absorption intensity. Notably, these low energy gaps are beneficial for strong absorption in the NIR biological window.

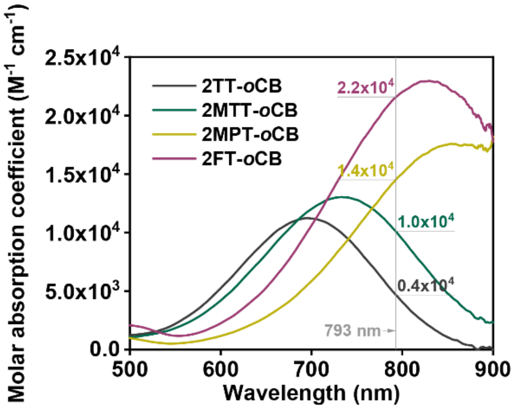


**Supplementary Fig. 17**. The spectra of the four AIEgens in THF showing the molar absorption coefficients at various wavelengths.

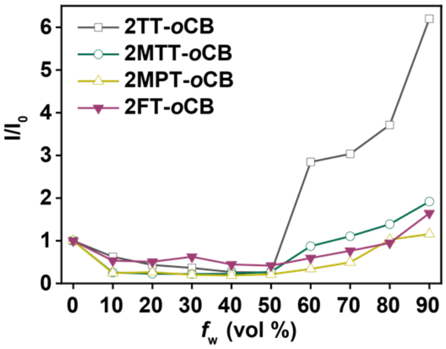


**Supplementary Fig. 18**. The plot of the PL peak intensity of the molecules versus $f_w$. $I$ and $I_0$ represent the peak intensities in the mixture with specific $f_w$, and the pure THF ($f_w = 0$).

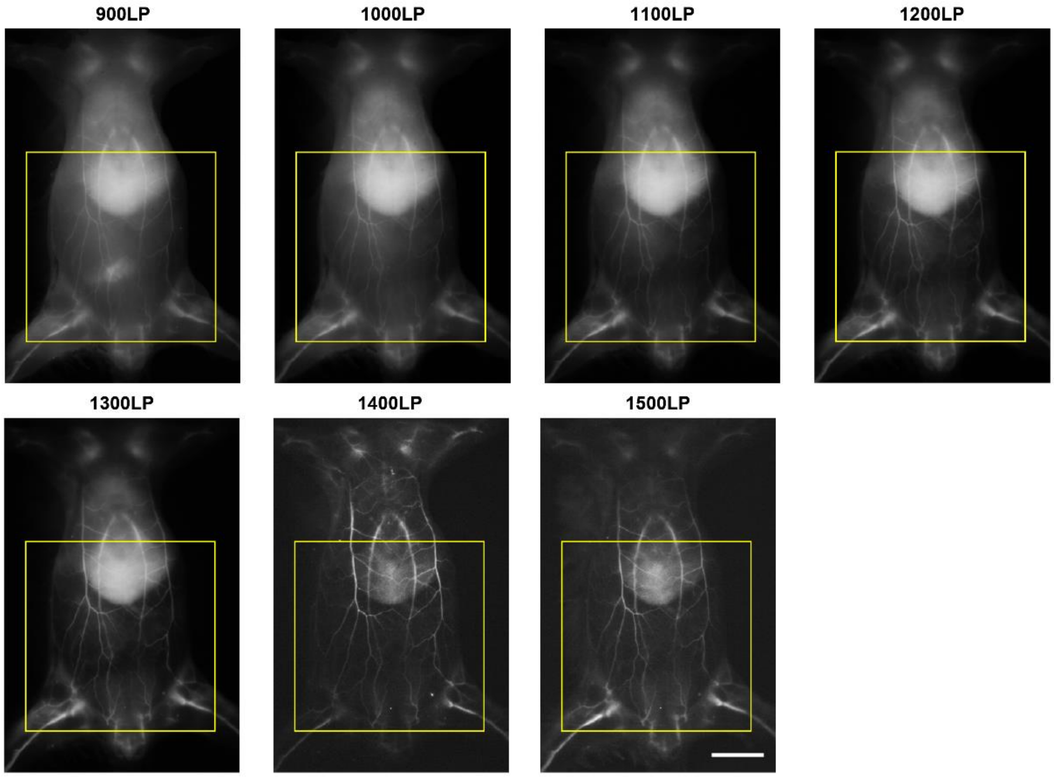


**Supplementary Fig. 19**. The NIR-II fluorescence whole-body vessel imaging in a mouse in different spectral regions. The selected areas with yellow squares were the original images of the FFT results in Supplementary Fig. 20. Scale bar, 10 mm.

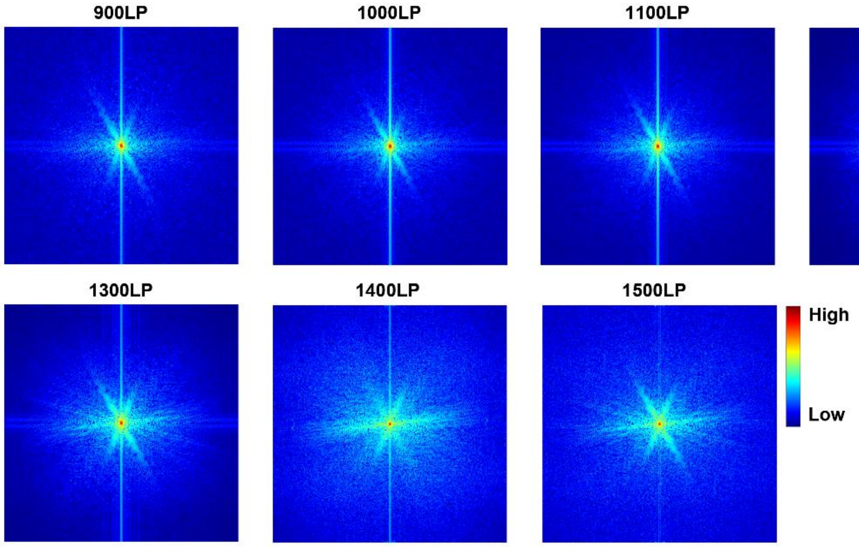

**Supplementary Fig. 20**. The FFT results of the NIR-II fluorescence vessel images in Supplementary Fig. 19. The spatial frequency gradually increases outward from the center of the map, and the color bar indicates the intensity.

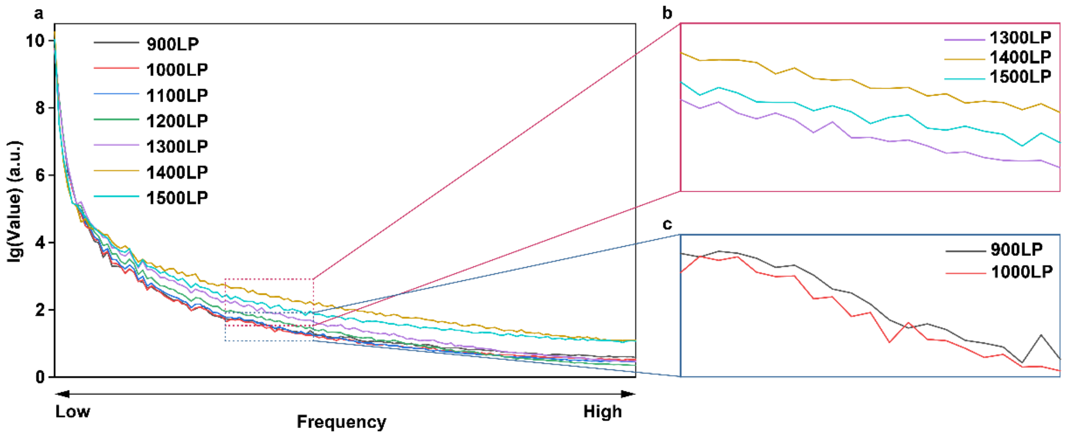


**Supplementary Fig. 21**. The frequency distributions of the FFT results. (a) The frequency distributions with 900LP, 1000LP, 1100LP, 1200LP, 1300LP, 1400LP, and 1500LP detection. (b) The enlarged region for direct comparison of 1300LP, 1400LP, and 1500LP imaging. (c) The enlarged region for direct comparison of 900LP and 1000LP imaging.

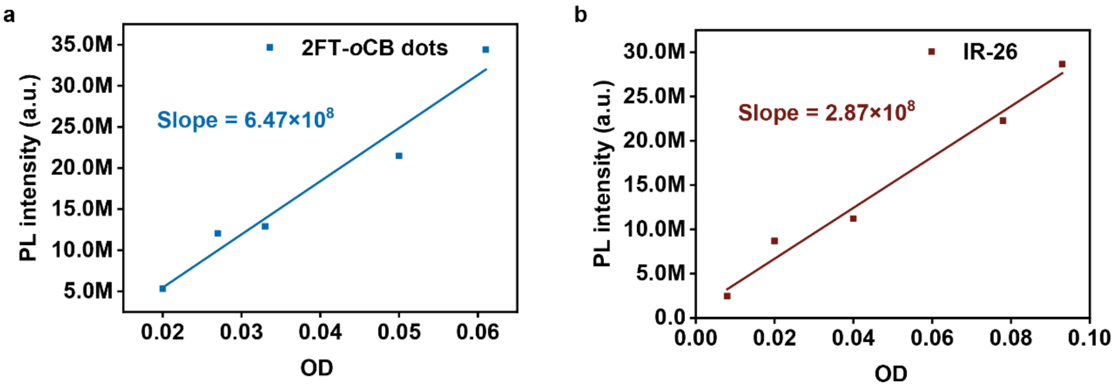


**Supplementary Fig. 22**. The measurement of the quantum yield beyond 1000 nm. The plots for the integrated fluorescence intensities of (a) 2FT-*o*CB dots in heavy water and (b) IR-26 in the 1, 2-dichloroethane at five different concentrations.

The integrated emission intensities of the 2FT-*o*CB dots and IR-26 were recorded under identical excitation conditions. The intensities were plotted as a function of OD, respectively, and the slopes were calculated by linear fittings. The quantum yield beyond 1000 nm of the sample was calculated as follows:

$$QY = QY_{ref} \cdot \frac{Slope}{Slope_{ref}} \cdot \frac{n^2}{n_{ref}^2}$$

Where $QY$ is the quantum yield of sample or reference, $n$ is the index of refraction of the solvent, and *Slope* is the slope value in Supplementary Fig. 22. Subscripts *ref* identify the reference.

Then, the QY beyond 1400 nm was calculated as follows:

$$QY_{>1400\ nm} = \frac{N_{emit>1400\ nm}}{N_{abs}} = \frac{N_{emit>1000\ nm}}{N_{abs}} \times \frac{N_{emit>1400\ nm}}{N_{emit>1000\ nm}} = QY_{>1000\ nm} \times \frac{I_{>1400\ nm}}{I_{>1000\ nm}}$$

Where $I_{>1400\ nm}$ *and* $I_{>1000\ nm}$ are the integrated fluorescence intensities of 2FT-*o*CB dots beyond 1400 nm and 1000 nm, respectively, and $\frac{I_{>1400\ nm}}{I_{>1000\ nm}}$ of 2FT-*o*CB dots could be calculated from the PL spectrum shown in Fig. 2g.

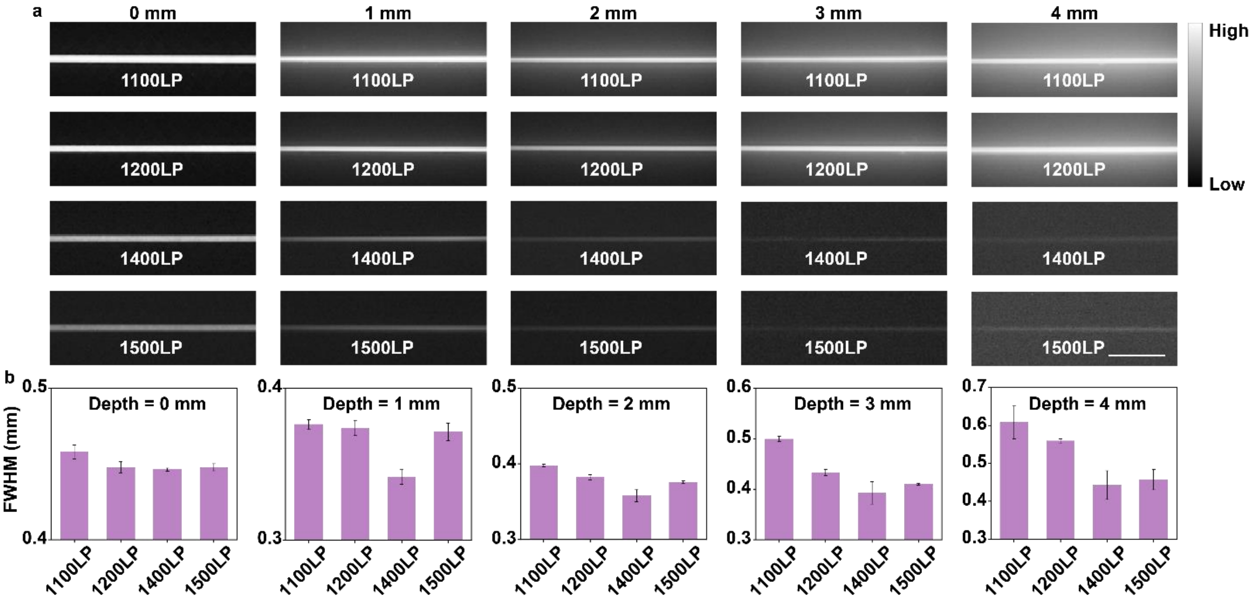


**Supplementary Fig. 23**. The NIR-II fluorescence phantom imaging in different spectral regions. **(a)** The phantom images of capillaries filled with the deuterium oxide dispersion of 2FT-*o*CB dots at depths of 0, 1, 2, 3, and 4 mm in 1% Intralipid® solution with 1100-, 1200-, 1400-, 1500-nm LP. Scale bar, 5 mm. **(b)** The FWHM measurements of the capillaries in Fig. 2a. Error bars indicate s.e.m. (n = 3). Scale bar, 5 mm.

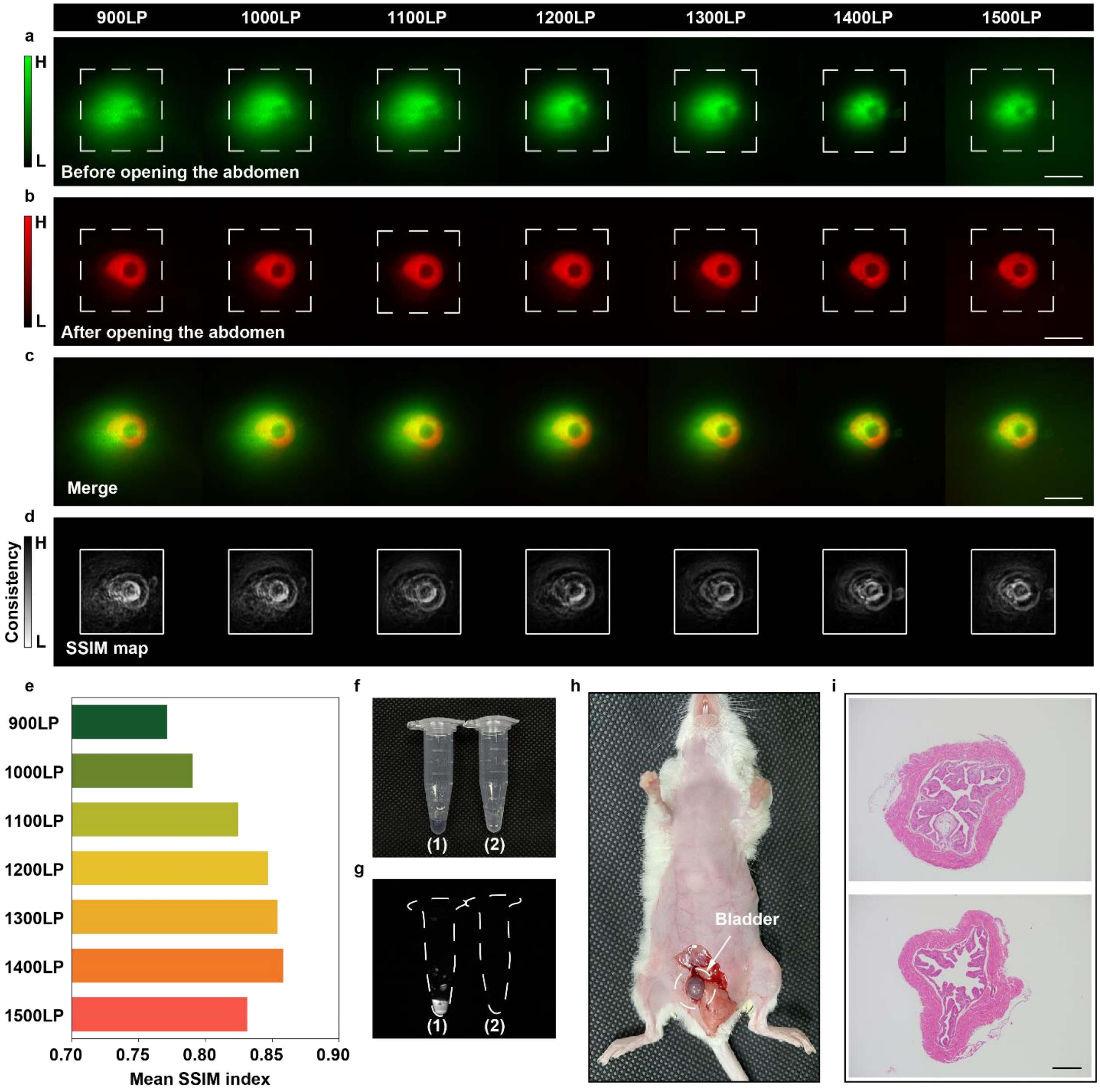


**Supplementary Fig. 24**. **The bladder fluorescence imaging in a mouse with high consistency.** The images of fluorescence cystography with intravesical instillation of the 2FT-oCB dots **(a)** before as well as **(b)** after opening the abdomen. **(c)** The merged images processed via image J. **(d)** The SSIM maps output from MATLAB. Scale bars, 5 mm. **(e)** The SSIM index of the seven SSIM maps in different collection spectral regions. The **(f)** bright-field and **(g)** NIR-II fluorescence images of the two tubes containing urine and 1×PBS, respectively. **(h)** The picture of the mouse with the filling of the bladder after opening the abdomen. **(i)** Representative histological images of the bladders of 1×PBS-treated (upper) and dots-treated (bottom) mice.

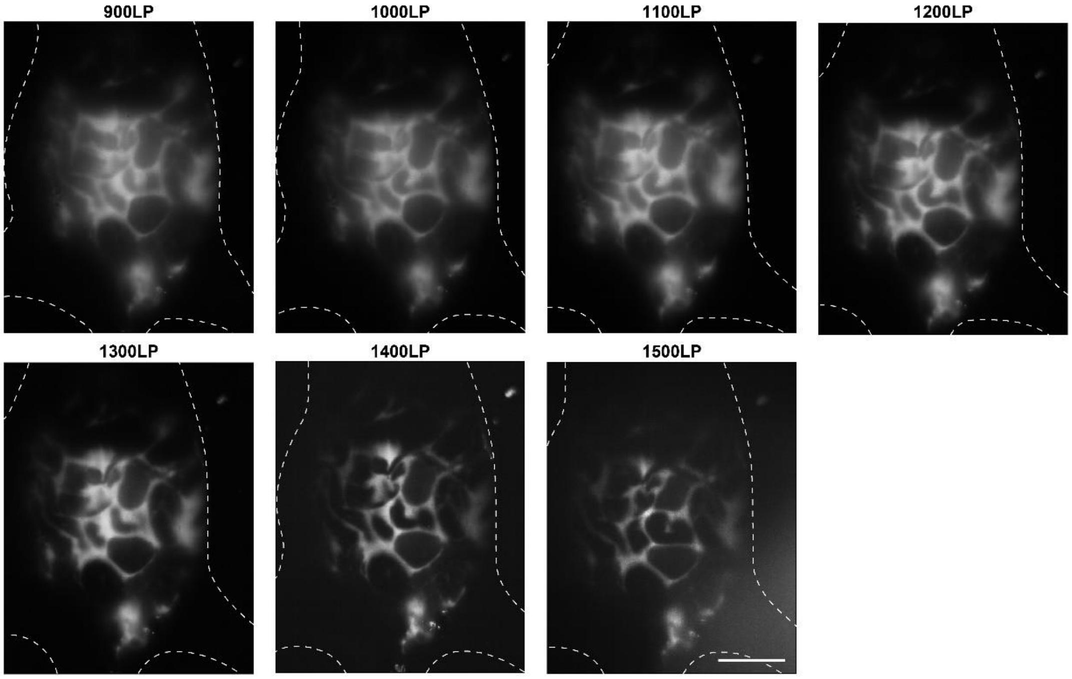


**Supplementary Fig. 25**. The whole-body imaging of a mouse with intraoperative bladder injury in different collection spectral regions. Scale bar, 10 mm.

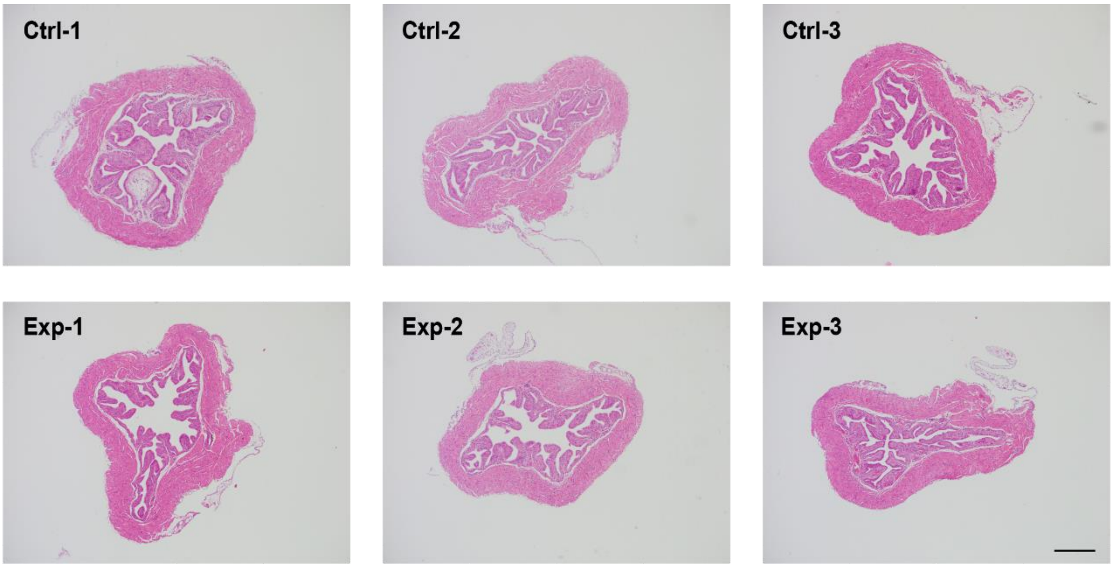


**Supplementary Fig. 26**. The H&E staining analysis of the bladders from mice after vesical perfusion with 1×PBS (Control group: Ctrl-1, Ctrl-2 and Ctrl-3; n = 3) and 2FT-*o*CB dots (Experimental group: Exp-1, Exp-2 and Exp-3; n = 3). Scale bar, 500 μm.

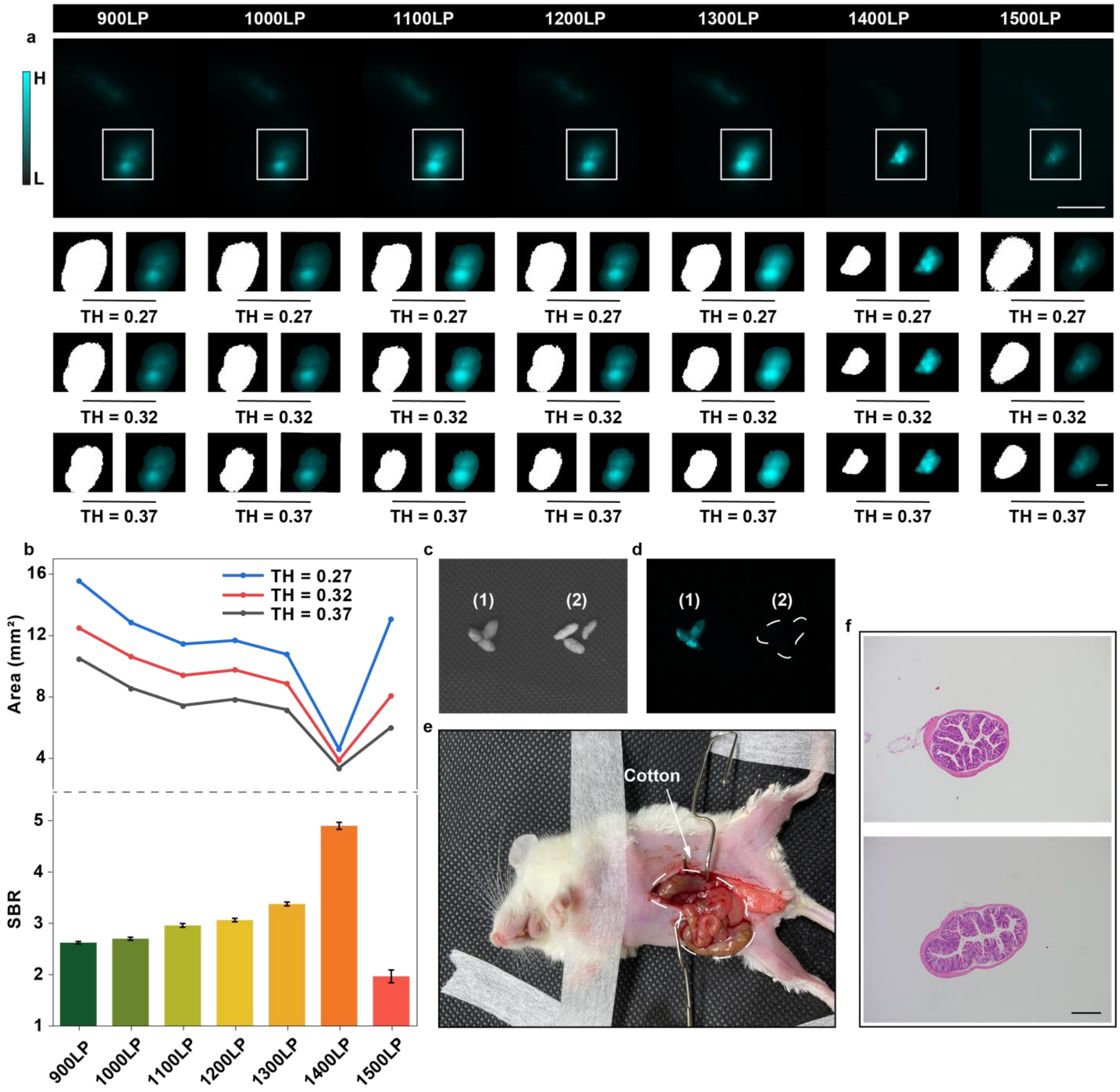


**Supplementary Fig. 27**. **The cotton fluorescence imaging in a mouse with high contrast. (a)** The images of fluorescence colonography (Scale bar, 5 mm) with colonic perfusion of 2FT-oCB dots, and the binary and segmented images (Scale bar, 1 mm) with the TH of 0.27, 0.32 and 0.37. **(b)** The signal areas after binarization in different collection spectral regions and the SBRs in different collection spectral regions. Error bars indicate s.e.m. (n = 3). The **(c)** bright-field and **(d)** NIR-II fluorescence images of the feces from the mouse treated with (1) dots and (2) 1×PBS. **(e)** The picture of the mouse with the filling of the colon after opening the abdomen. **(f)** Representative histological images of the colons of 1×PBS-treated (upper) and dots-treated (bottom) mice.

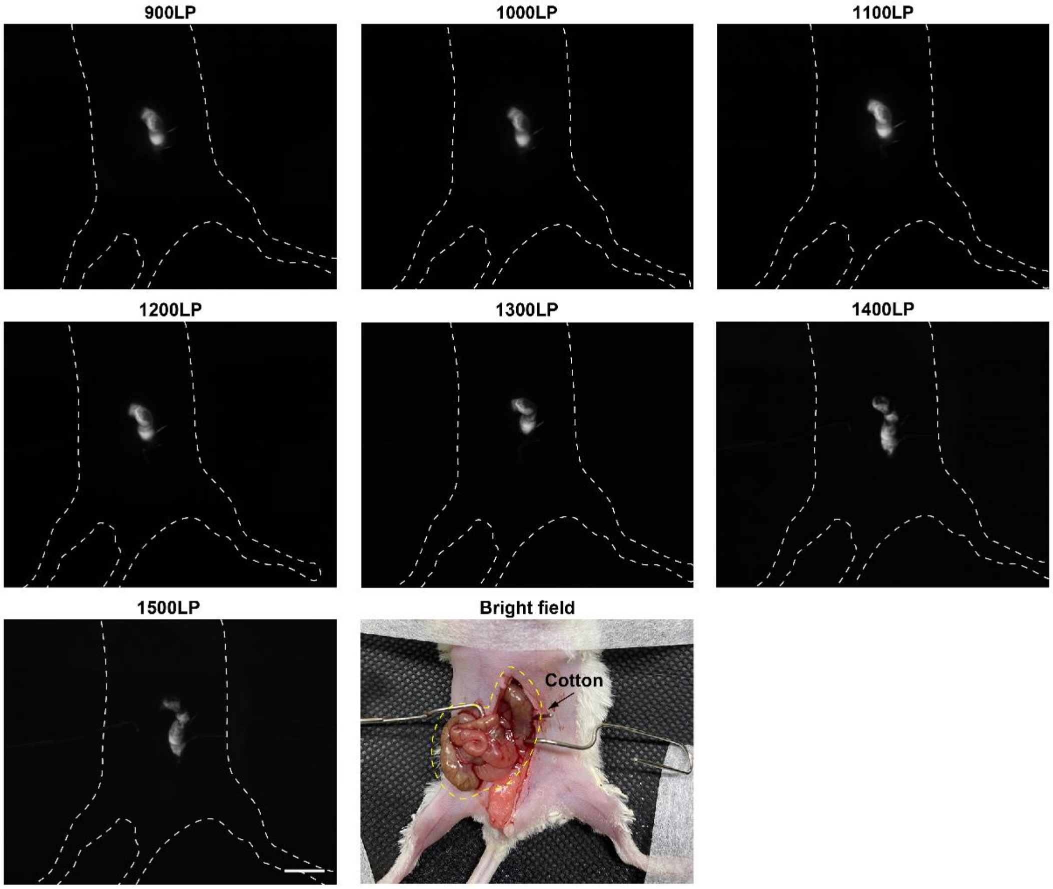


**Supplementary Fig. 28**. The NIR-II fluorescence colon images after opening the abdomen in different collection spectral regions. Scale bar, 5 mm.

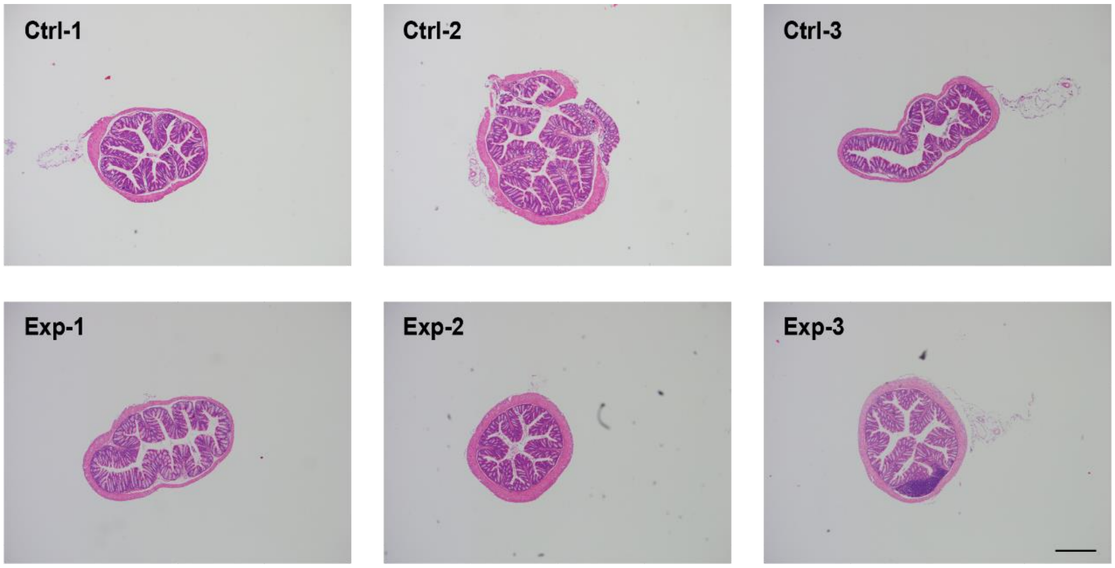

**Supplementary Fig. 29**. The H&E staining analysis of the colons from mice after colonic perfusion with 1×PBS (Control group: Ctrl-1, Ctrl-2 and Ctrl-3; n = 3) and 2FT-*o*CB dots (Experimental group: Exp-1, Exp-2 and Exp-3; n = 3). Scale bar, 500 μm.

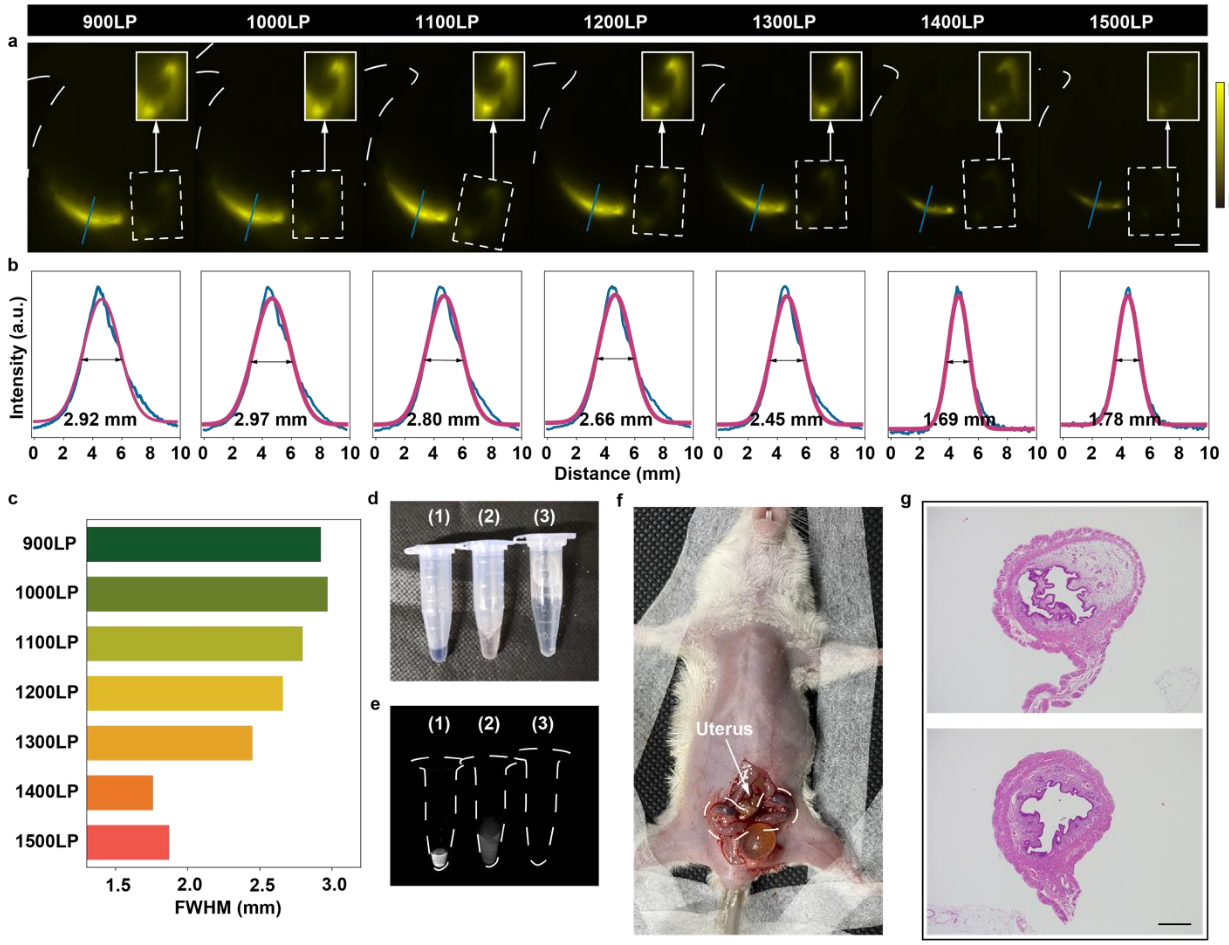


**Supplementary Fig. 30**. **The uterus fluorescence imaging in a mouse with high spatial resolution. (a)** The images of fluorescence hysterography with uterine perfusion of 2FT-oCB dots. Scale bar, 5 mm. The inserts are enhanced right side. Scale bar, 3 mm. **(b)** Cross-sectional fluorescence intensity profiles along the blue lines. The white dashed lines in Supplementary Fig. 30a describe the outline of the mouse body. **(c)** The FWHMs in different collection spectral regions. The **(d)** bright-field and **(e)** NIR-II fluorescence images of the three tubes containing the (1) liquid flowing out naturally from the uteruses, (2) intrauterine lavage fluid, and (3) 1×PBS, respectively. **(f)** The picture of the mouse with the filling of the uterus after opening the abdomen. **(g)** Representative histological images of the uteruses of 1×PBS-treated (upper) and dots-treated (bottom) mice.

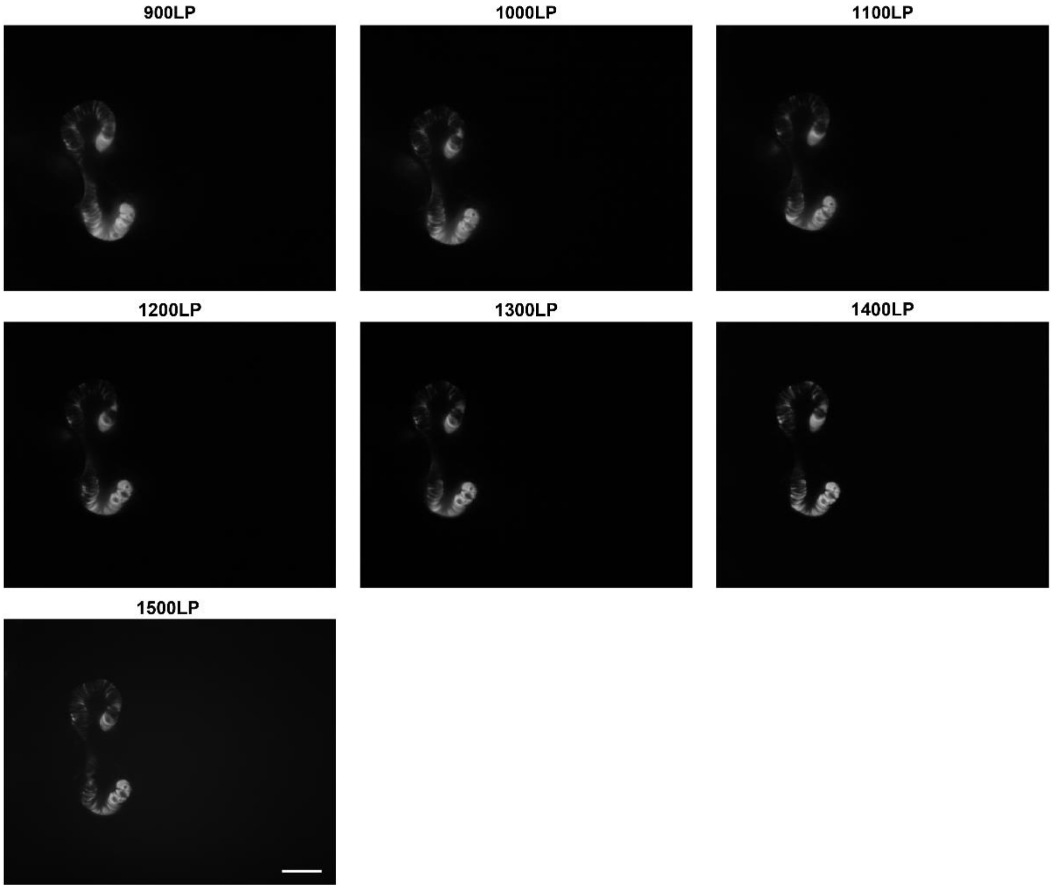


**Supplementary Fig. 31**. The NIR-II fluorescence uteruses images after opening the abdomen in different collection spectral regions. Scale bar, 5 mm.

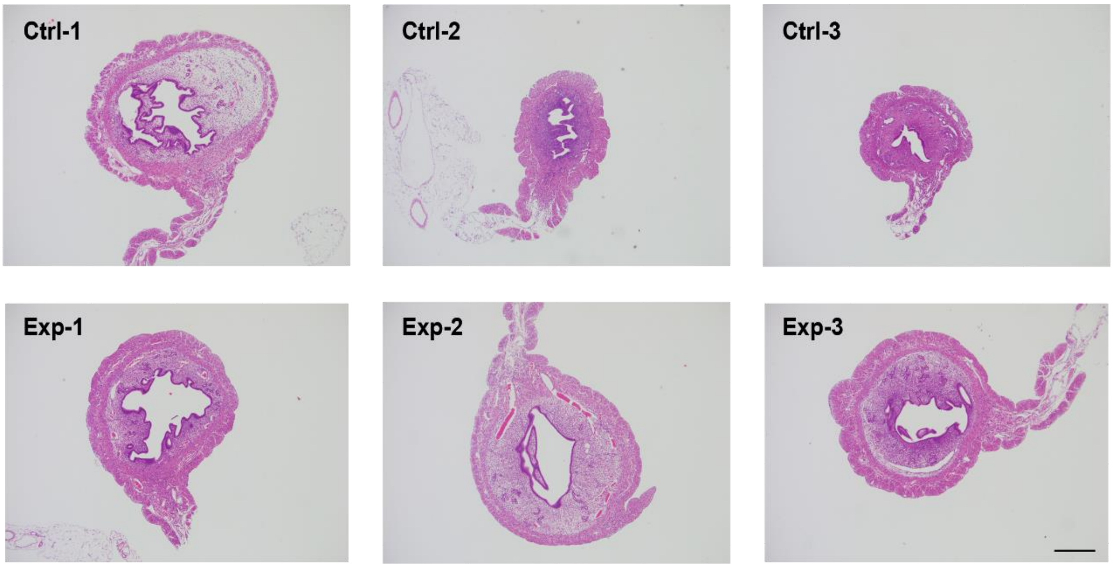

**Supplementary Fig. 32**. The H&E staining analysis of the uteruses from mice after uterine perfusion with 1×PBS (Control group: Ctrl-1, Ctrl-2 and Ctrl-3; n = 3) and 2FT-*o*CB dots (Experimental group: Exp-1, Exp-2 and Exp-3; n = 3). Scale bar, 500 μm.

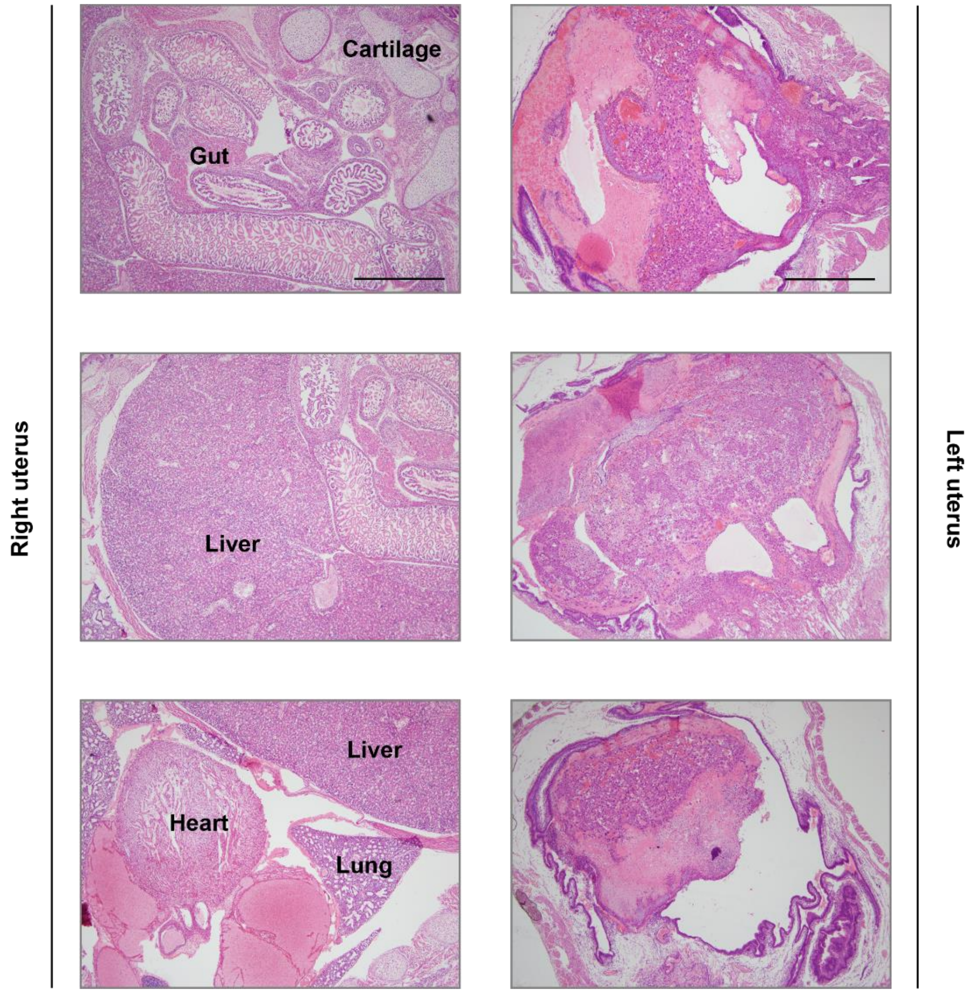


**Supplementary Fig. 33** The H&E staining analysis of the fetus in the right uterus and the residual pregnancy tissue in the left uterus from mice. Scale bars, 500 μm.

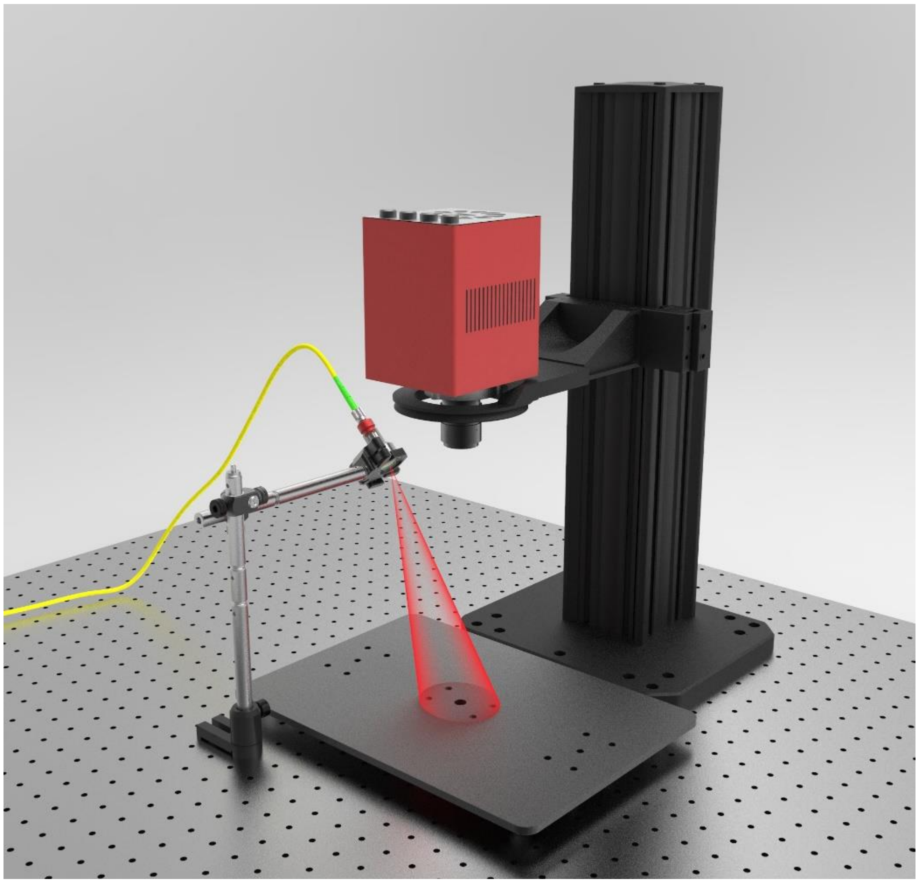

**Supplementary Fig. 34**. NIR-II fluorescence macro imaging system.

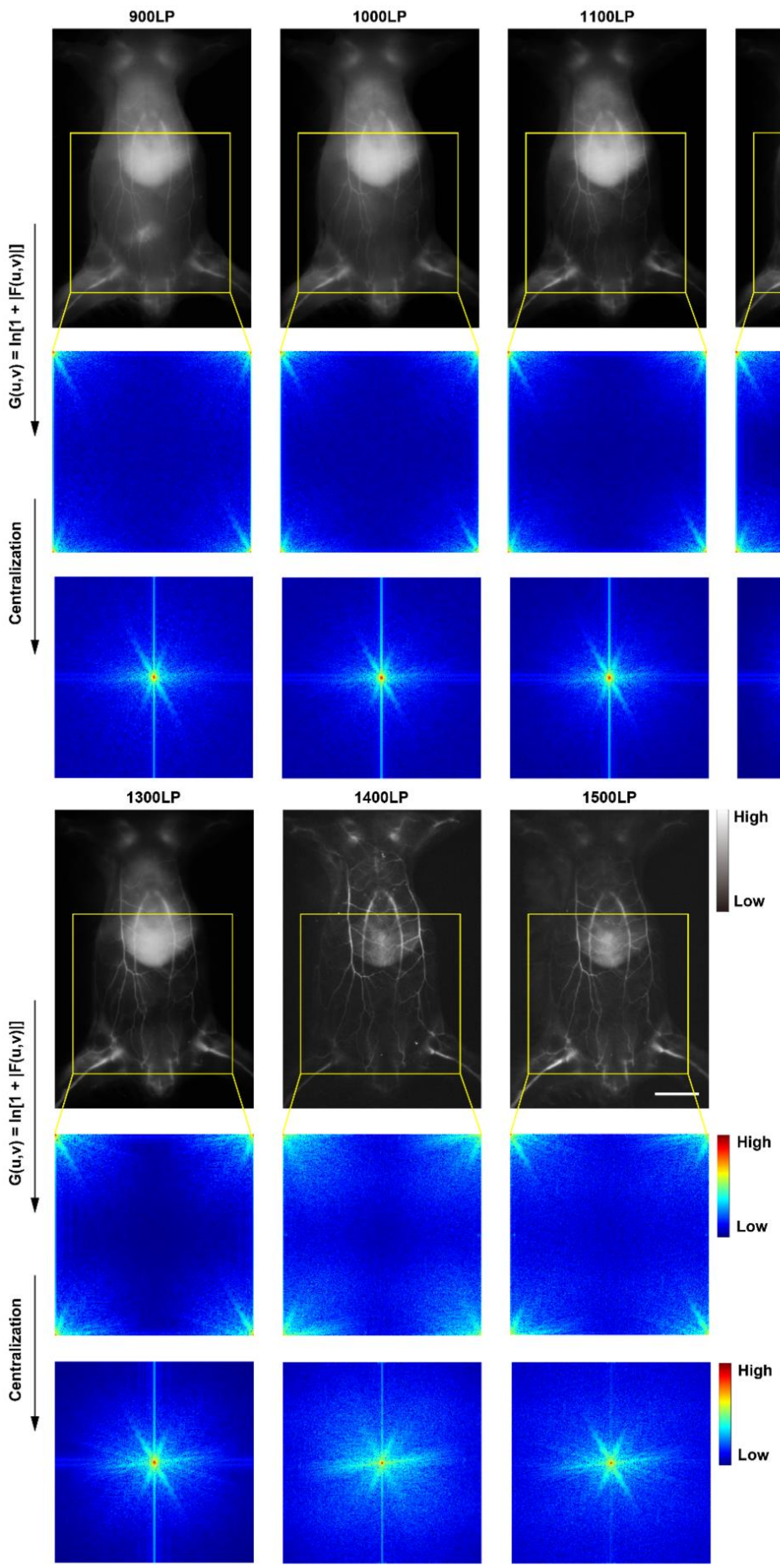


**Supplementary Fig. 35**. The whole image processing of the FFT results. Scale bar, 10 mm.

**Supplementary Table 1**. The power densities and the exposure times in the experiments.

| Long-pass detection (LP) | **Whole-body vessel imaging** | | **Fluorescence cystography** | | **Fluorescence hysterography** | | **Fluorescence colonography** | |
|---|---|---|---|---|---|---|---|---|
| | Power density (mW/cm$^2$) | Exposure time (ms) | Power density (mW/cm$^2$) | Exposure time (ms) | Power density (mW/cm$^2$) | Exposure time (ms) | Power density (mW/cm$^2$) | Exposure time (ms) |
| 900 | ~20 | 50 | ~10 | 50 | ~15 | 50 | ~20 | 50 |
| 1000 | ~20 | 50 | ~10 | 50 | ~15 | 50 | ~20 | 50 |
| 1100 | ~40 | 50 | ~10 | 80 | ~20 | 100 | ~40 | 50 |
| 1200 | ~40 | 200 | ~20 | 80 | ~40 | 100 | ~40 | 100 |
| 1300 | ~100 | 500 | ~60 | 100 | ~60 | 200 | ~80 | 200 |
| 1400 | ~120 | 800 | ~100 | 500 | ~120 | 500 | ~160 | 500 |
| 1500 | ~120 | 1000 | ~100 | 500 | ~120 | 500 | ~160 | 500 |
